# Temporal constraints on enhancer usage shape the regulation of limb gene transcription

**DOI:** 10.1101/2024.03.22.585864

**Authors:** Raquel Rouco, Antonella Rauseo, Guillaume Sapin, Olimpia Bompadre, Fabrice Darbellay, Guillaume Andrey

## Abstract

Repertoires of transcriptional enhancers orchestrate gene expression during embryonic development, thereby shaping the forms and functions of organs. Within these repertoires individual enhancers display spatially distinct or overlapping activities that collectively build up the expression domain of cognate genes. However, the temporal specificity of these enhancers - how their activities change over developmental time to dynamically influence gene expression - remains uncharacterized. Here, we observed that temporally restricted enhancer repertoires are embedded at numerous loci associated with mouse limb development. To monitor how such enhancer repertoires govern gene transcription *in vivo* across extensive developmental periods, we introduce the regulatory trajectory framework. This paradigm conceptually involves transcriptional initiation, marking the beginning of gene expression, followed by its maintenance over time, and ultimately decommissioning, leading to gene repression. To track and sort cells undergoing these distinct phases, we devised a transgenic recorder approach at the *Shox2* model locus. Through this method, we discovered that cells maintaining *Shox2* transcription in early and late limb development relies on distinct, temporally restricted enhancer repertoires. We demonstrate that eliminating early-or late-acting enhancers only transiently affects *Shox2* expression indicating that these enhancer repertoires function independently. Additionally, we found that changes in the 3D topology of the locus associate with enhancer activities and that a rapid loss of enhancer-promoter contacts occurs during decommissioning. Finally, we show that the decommissioning of the *Shox2* locus can be actively driven by *Hoxd13*, a gene which expression is known to antagonize *Shox2*. Overall, our work uncovers the dependency of developmental genes on enhancers with temporally restricted activities to generate complex expression patterns over time and shed light on the dynamics of enhancer-promoter interactions.

## Introduction

Organ and tissue patterning depend on spatiotemporally defined and cell-specific transcriptional activities, which are regulated by enhancers, among other regulatory elements (Petit et al., 2017; Robson et al., 2019). Enhancers bear the ability to be bound by transcription factors integrating the cellular environments and ultimately to translate it into an integrated transcriptional output (Spitz and Furlong, 2012). Within the same chromatin domains known as Topologically Associating Domains (TADs), genes can dynamically interact with multiple enhancers, referred to as enhancer repertoires, collectively shaping their expression patterns (Abassah-Oppong et al., 2023; Andrey and Mundlos, 2017; Furlong and Levine, 2018; Kvon et al., 2021; Robson et al., 2019; Stadhouders et al., 2019). While multiple studies have explored how distinct enhancers may act with distinct or overlapping spatial activities (Frankel et al., 2010; Kvon et al., 2014; Osterwalder et al., 2018; Perry et al., 2010; Petit et al., 2017; Robson et al., 2019; Werner et al., 2007), their temporal specificities remain less clear. In the context of a growing and differentiating embryo, it is unknown how regulatory landscapes adapt to the changing *trans*-and signaling environments to sustain target gene transcription over time. It is probable that enhancer repertoires change over time; however, the independence of these repertoires in establishing the definitive expression pattern of genes has yet to be fully understood. Furthermore, although several groups have investigated how enhancers are decommissioned, the way large regulatory landscapes, involving multiple enhancers, terminate their activities is still not well described (Respuela et al., 2016; Whyte et al., 2012; Wu et al., 2023).

Many complex regulatory landscapes have been dissected in the limb model system (Kragesteen et al., 2018; Malkmus et al., 2021; Petit et al., 2017; Will et al., 2017). The mouse limb bud is a complex appendage formed by different cell types that constitutes a widely used organ to study development. Fore- and hindlimbs are budding from the lateral plate mesoderm at E9.5 and E10.0 respectively (Martin, 1990). Initially, the limb is predominantly composed of undifferentiated mesenchyme, capable of forming all proximo-distal segments. However, as development progresses, this potential gradually diminishes as cells set their proximal or distal identity, and differentiate into cartilage and connective tissues (Cooper et al., 2011). Mechanistically, as limb progenitor cells begin expressing patterning genes, they can either commit to the associated specific limb segment or maintain their progenitor status by activating more distal factors and repressing the more proximal ones. Thus, once a proximo-distal patterning gene is activated in limb progenitors, its expression can either be maintained during lineage commitment or inhibited to permit differentiation into a different, more distal limb segment (Markman et al., 2023; Saiz-Lopez et al., 2015; Verheyden and Sun, 2008). Therefore, the limb is an outstanding model system to define how regulation can be sustained or repressed over time.

In this study, we sought to characterize the regulatory mechanisms of transcriptional maintenance over time and locus-wide decommissioning within the limb developmental context. To allow for the functional dissection of these mechanisms, we are using the *Shox2* gene locus that bear an essential role in proximal limb development. *Shox2* expression starts in the limb around E9.5/E10.0 and continues to be expressed in proximal connective tissues and cartilage, playing a crucial role in the growth of proximal limb bones, such as the humerus and the femur (Blaschke et al., 2007; Cobb et al., 2006; Espinoza-Lewis et al., 2009; Glaser et al., 2014; Gu et al., 2008; Neufeld et al., 2014; Rosin et al., 2015; Scott et al., 2011; Yu et al., 2005). Moreover, the severe shortening of proximal limb bone segments which has been observed in conditional knockout approaches (Blaschke et al., 2007; Glaser et al., 2014; Yu et al., 2007) serves as a good model for human short stature caused by mutations in the *SHOX* gene, a paralog absent in rodents (Clement-Jones et al., 2000; Cobb et al., 2006; Gianfrancesco et al., 2001).

The regulatory landscape of *Shox2* is embedded within a 1.1 Mb TAD that splits into a centromeric side featuring a 500 kb gene desert and a telomeric side, largely composed of the introns of the *Rsrc1* gene. Numerous limb enhancers have been identified on both the telomeric and centromeric sides, collectively contributing to *Shox2* limb expression (Abassah-Oppong et al., 2023; Osterwalder et al., 2018; Rosin et al., 2013; Ye et al., 2016). Although these enhancer regions have been shown to drive expression in the limb and other *Shox2*-expressing tissues, the specific time windows of their activity have not been determined.

Given that *Shox2* expression is initiated in the early, undifferentiated limb bud and gradually becomes restricted to proximal limb mesodermal derivatives, its regulation must involve mechanisms of transcriptional maintenance and repression *via* precise enhancer activation and decommissioning, respectively. To accurately track descendants of *Shox2*-expressing cells within the complex and diversely populated developing limb, we have devised a novel conceptual and experimental framework to follow *Shox2* locus activity. This approach enables the isolation of cells at distinct phases of *Shox2* regulation, offering new insights into the dynamic control of gene expression during limb development.

## Results

### Characterization of temporal enhancer repertoire changes at limb development associated *loci*

To examine whether *Shox2*, and more generally limb developmental genes, could rely on distinct enhancer repertoire over time during limb development, we first re-analyzed transcriptomic and epigenomic data from the E10.5 and E13.5 limb (Andrey et al., 2017). Specifically, we selected 90 genes relevant for limb development, including *Shox2*, whose expression levels remained stable between both stages. We mapped for these genes putative enhancers using H3K27ac ChIP-seq enrichment within their contact domains, as defined by promoter Capture-C in the limb (Andrey et al., 2017; Rada-Iglesias et al., 2011) (**Supplementary Fig. S1A, Supplementary Table S1**).

Across the 90 investigated contact domains, we observed a total of 1,626 putative enhancers. Of these, 402 were active at both E10.5 and E13.5 (25%, termed common enhancers), 506 were specifically active at E10.5 (31%, termed early enhancers), and 718 at E13.5 (44%, termed late enhancers) (**Fig. 1A, Supplementary Table S1**). A large majority of these contact domains (76 of them, 84%), including *Shox2*, contained all three types of enhancers, a smaller fraction (13%) presents at least two types while only two loci display putative enhancers belonging to a unique category (**Fig. 1A, Supplementary Table S1**). This observation, which is exemplified in **Fig. 1B** by *Shox2* and two well-known limb-associated loci, *Sox11* and *Sox9,* suggests that within the limb context, the usage of putative enhancer repertoires generally shift over time (**Supplementary Fig. S1A**). This analysis led us to conclude that the *Shox2* locus, which display 13 early, 3 common and 4 late putative enhancers (**Fig 1B, Supplementary Fig. S1A**), is representative of limb developmental genes to study how genetic regulations are controlled over time.

**Figure 1:**
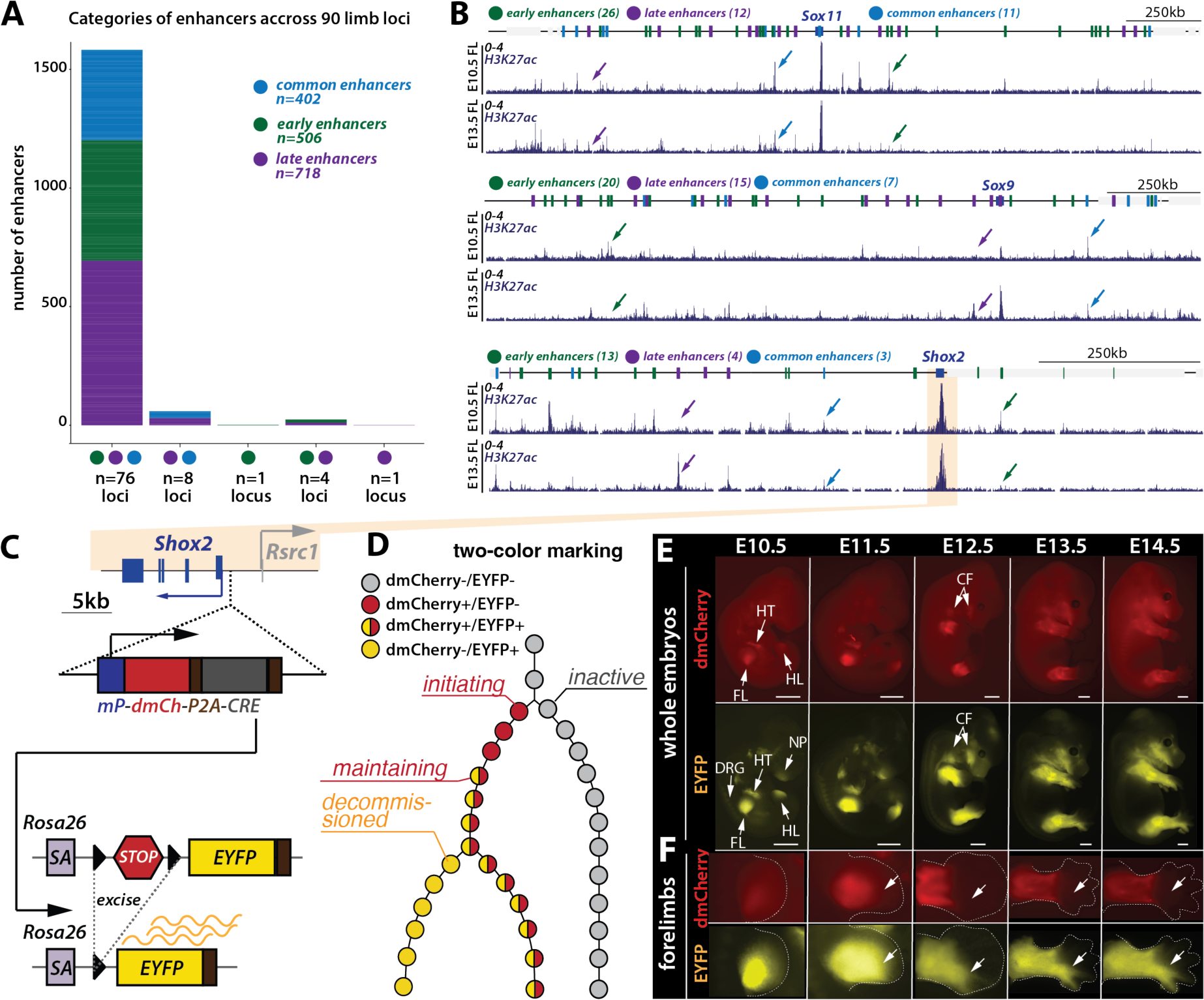
Temporal usage of enhancers repertoire hypothesis studied by a dual-color reporter at Shox2. **A.** Distribution of 90 limb-related loci based on the presence of putative early (green), common (light blue), and late (purple) enhancers defined by H3K27ac enrichment on forelimbs at E10.5 and E13.5 within their contact domain. **B.** H3K27ac ChIPseq tracks at from E10.5 and E13.5 forelimbs of three examples of developmental loci: Sox11 contact domain shown, mm39; chr12:26470941-28552347), Sox9 (contact domain shown, mm39; chr11:111354041-113105934) and Shox2 (contact domain shown, mm39; chr3:66194862-67308968). Early (green spheres), common (blue spheres), and late (purple spheres) putative enhancer regions marked on the locus illustration are based on H3K27Ac ChIP profile re-analyzed from (Andrey et al., 2017). Light grey box represents other genes. For each locus, arrows pinpoint a representative example of one early (green), one common (blue) and one late (purple) putative enhancers. **C.** A double fluorophore approach to monitor Shox2 locus activity over time; mP: minimal β-globin promoter, dmCh: mCherry gene with a destabilized PEST sequence; P2A: self-cleavage sequence; CRE: CRE recombinase gene; SA: splice acceptor; STOP: floxed 3xSV40pA STOP signal; EYFP: EYFP gene. **D.** Schematic of the double fluorophore approach that enables tracking of Shox2 regulatory trajectory: different color combinations correlate with different phases of the trajectory. **E.** Imaging of dmCherry and EYFP fluorescence in Shox2^trac^ (Shox2^dmCherry/+^; Rosa26^loxEYFP/+^) embryos (scale: 1mm); HT: heart, FL: forelimbs, HL: hindlimbs, NP: nasal process, DRG: dorsal root ganglia, and CF: craniofacial structures. **F.** Imaging of dmCherry and EYFP fluorescence in developing Shox2^trac^ forelimbs. Note that cells in digits 4 and 5 (white arrows) are positive for EYFP but negative for dmCherry.

### A transgenic reporter system to characterize the *Shox2* regulatory trajectory

To carefully analyze the regulatory dynamics of gene expression across extended developmental periods, we conceptually divide it into different states that collectively define a regulatory trajectory. Such a trajectory begins from a poised state, from which a locus may transit towards an active state *via* transcriptional *initiation*, or towards an inactive state *via repression* (**Supplementary Fig. S1B**) (Bernstein et al., 2006; Mas et al., 2018; Mikkelsen et al., 2007). After transcriptional *initiation*, it is eventually followed by a *maintenance* phase where gene transcription is sustained over time. Finally, a *decommissioning* phase progressively leads to gene repression (**Supplementary Fig. S1B**).

To monitor the *Shox2* regulatory trajectory and sort cells undergoing these distinct phases, we have developed a dual-color transgenic reporter system. First, we inserted in mouse embryonic stem cells (mESCs), 1kb upstream of the *Shox2* transcriptional start site, a regulatory sensor cassette constituted by a minimal *β-globin* promoter, a mCherry reporter open reading frame (ORF) followed by a destabilized PEST sequence, a P2A self-cleavage sequence and the CRE recombinase ORF (*Shox2^dmCherry^*^/+^) (**Fig. 1C**)(Akhtar et al., 2013; Kondo and Duboule, 1999; Marinic et al., 2013; Ruf et al., 2011; Symmons et al., 2016). These *Shox2^dmCherry/+^* cells were then retargeted to integrate, at the *Rosa26* locus, a cassette with a splice acceptor followed by a floxed 3x SV40pA STOP signal and the EYFP ORF (*Shox2^dmCherry/+^*; *Rosa26^loxEYFP/+^ or Shox2^trac^*) (**Fig. 1C**) (Srinivas et al., 2001). With this system, *Shox2* expressing cells will induce dmCherry-P2A-CRE transcription and also continuously express EYFP, even after the inactivation of *Shox2* and dmCherry-CRE transcription. Here, cells undergoing transcriptional *initiation* should correspond to early limb *Shox2* expression and dmCherry-positive cells (**Fig. 1D**). Cells with dmCherry and EYFP signals will indicate transcriptional *maintenance* and cells with EYFP only will have undergone transcriptional *decommissioning* after an earlier active transcriptional phase, while cells that remained inactive will have no fluorescent labelling (**Fig. 1D**).

Embryos were then obtained from these *Shox2^trac^* mESCs by tetraploid complementation (Artus and Hadjantonakis, 2011). At E10.5, we detected dmCherry and EYFP signals in the fore-(FL) and hindlimbs (HL) as well as in the heart (HT) (**Fig 1E**). We also detected EYFP signal in the nasal process (NP) and dorsal root ganglia (DRG). While the dmCherry signal in hindlimbs was weak at E10.5 compared to forelimbs, it markedly increased by E11.5, reflecting the expected developmental delay of hindlimbs compared to forelimbs (Martin, 1990). Subsequently, at E12.5, additional craniofacial (CF) structures exhibited fluorescence from both dmCherry and EYFP (**Fig 1E**). These expression patterns closely match the known expression profile of *Shox2* (**Supplementary Fig. S1C, D**) (Blaschke et al., 1998; Glaser et al., 2014; Semina et al., 1998; Sun et al., 2013). Intriguingly, within the limbs, we observed that digits 4 and 5, located in the posterior autopod, were exclusively marked by EYFP and not by dmCherry (**Fig. 1E, F, Supplementary Fig. S1D**). This indicates that part of the distal posterior limb originates from progenitor cells that had expressed *Shox2* at earlier stages, but whose regulatory landscape was subsequently decommissioned.

### Single-cell insights into *Shox2* transcriptional dynamics trace early *decommissioned* cells in the distal limb

To gain a detailed understanding of transcriptional dynamics during limb development, and of *Shox2* transcriptional phases over time, we produced single-cell RNA-seq transcriptomes from *Shox2^trac^* hindlimbs at four different embryonic stages: E10.5, E11.5, E12.5, and E13.5 (**Fig. 2A, Supplementary Fig. S2A**). We then specifically focus our analyses on mesenchymal cells (*Prrx1*+, *Prrx2*+, *Twist1*+), where *Shox2* is expressed (**Supplementary Fig. S2A, B**). Merging the four timepoints investigated together, we identified 15 distinct clusters (**Fig. 2B, Supplementary Table S2A**), from limb progenitors (**LP**, *Hoxd9/10/11*+, *Tfap2c*+, *Msx1*+, *Sall3/4*+) composed mainly by cells from E10.5 embryos, to more differentiated clusters at later stages (**Supplementary Fig. S2C-D**). To precisely delineate the developmental trajectories between these 15 clusters, we then ran a velocity analysis (Bergen et al., 2020; La Manno et al., 2018). We found that limb mesenchymal cells originate from the E10.5 limb progenitor cluster that is predicted to differentiate into both distal and proximal clusters (**Fig. 2B, C**). In proximal clusters, which are marked by *Shox2* expression (**Supplementary Fig. S3A, B**), progenitor pools (early and late proximal progenitors (**EPP** and **LPP**)) differentiate into proximal condensations (Proximal Growth Plate (**PGP**), Early and Late Proximal Condensations (**EPC** and **LPC**)) and connective tissues (Proximal Connective Tissue (**PCT**), Irregular Connective Tissue (**ICT**), and Tendon Progenitors (**TP**)). In distal clusters which are marked by *Hoxd13* expression (**Supplementary Fig. S3B**), progenitor pools (Early and Late Distal Progenitors (**EDP** and **LDP**)) differentiate into interdigit mesenchyme (**IM**) and distal condensations (Early and Late Digit Condensations (**EDC**, **LDC**)). The Mesopodium (**Ms**), the presumptive wrist domain, is predicted to derive directly from limb progenitors, expressing both proximal (*Shox2*) and distal markers (*Hoxd13*) (**Fig. 2B, C**).

**Figure 2:**
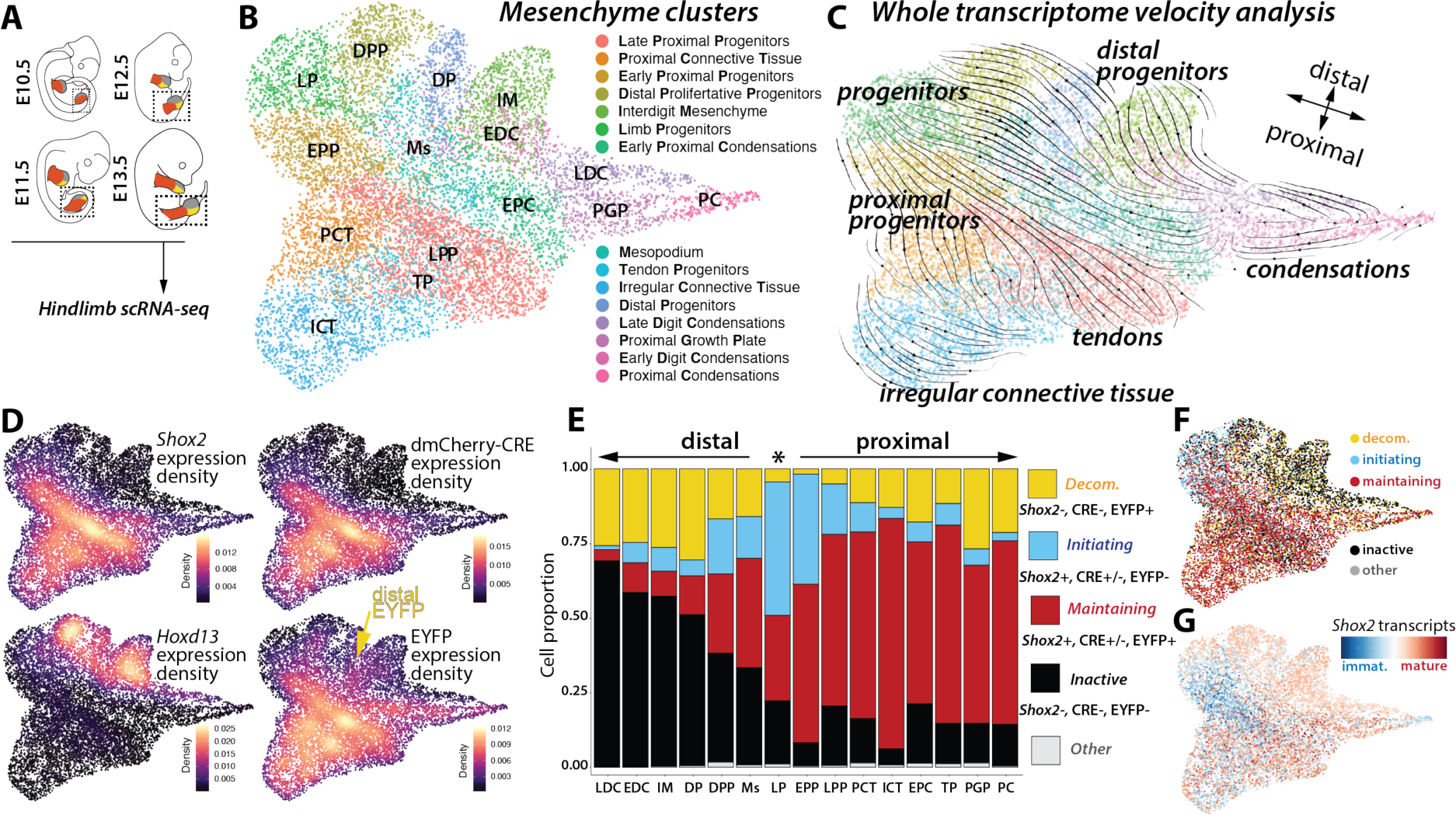
Single-cell analyses of Shox2 regulatory trajectory. **A**. Illustration of single-cell preparation from micro-dissected Shox2^trac^ hindlimbs at E10.5, E11.5, E12.5, and E13.5. **B**. UMAP visualization of re-clustered mesenchymal cells from all merged datasets. **C**. RNA-velocity analysis across mesenchymal cell clusters. **D**. Analysis of gene expression density for Shox2 (proximal marker), dmCherry-P2A-CRE, Hoxd13, and EYFP (distal marker) in mesenchyme cells. Note the EYFP expression in distal clusters, in contrast to Shox2 and dmCherry-CRE **E**. Distribution of cell proportions categorized by the different Shox2 transcriptional phases: initiation (Shox2+, dmCherry-P2A-CRE +/-, EYFP-), maintaining (Shox2+, dmCherry-P2A-CRE +/-, EYFP+), decommissioned (Shox2-, dmCherry-P2A-CRE-, EYFP+), inactive (Shox2-, dmCherry-P2A-CRE-, EYFP-) or other cells (when where not included in any of the previously mentioned class) across mesenchymal clusters. Clusters are ordered according to their distal and proximal identity and to developmental time, with the limb progenitor cluster highlighted by a star. **F**. UMAP representing different Shox2 cell phases: inactive (black), initiation (light blue), maintaining (red), decommissioned (yellow) and other (grey). **G**. Visualization of Shox2 RNA-velocity: light blue signifies a higher fraction of immature transcripts (immat.), whereas red indicates a higher fraction of mature transcripts.

We then mapped within this clustering *dmCherry-P2A-CRE* and *EYFP* expressing cells from E10.5 onwards (**Fig. 2D, Supplementary Fig. S2E, S3A**). As expected from imaging data (**Fig. 1E, Supplementary Fig. S1C**), we observed a high correlation between *Shox2* and *dmCherry-P2A-CRE* transcription (correlation coefficient=0.808, p=0.00024, where the p-value is the probability for the correlation coefficient to be negative (Lopez-Delisle and Delisle, 2022)) indicating that the dmCherry-P2A-CRE cassette recapitulates *Shox2* expression. Interestingly, *EYFP* expression was observed in both proximal and distal limb clusters, though less frequently in the latter (**Fig. 2D, Supplementary Fig. S3A**), matching with our previous imaging observations (**Fig. 1E, Supplementary Fig. S1D**). These data confirm that the *Shox2^trac^* system efficiently tracks *Shox2* transcriptional activity over time.

Leveraging the transcript levels of *Shox2*, EYFP, and dmCherry-P2A-CRE we then categorize cells into distinct phases of the *Shox2* regulatory trajectory (**Fig. 1D**). Given the low transcription levels of *dmCherry-P2A-CRE* in comparison to *EYFP* and *Shox2* (for technical reasons, see Material and Methods), we primarily depended on the latter two genes expression to annotate the locus activity, classifying cells as *inactive* (*Shox2-*, *dmCherry-P2A-CRE-, EYFP-*), *initiating* (*Shox2+*, *dmCherry-P2A-CRE +/-, EYFP-*), *maintaining* (*Shox2+*, *dmCherry-P2A-CRE +/-, EYFP+*), and *decommissioned* (*Shox2-*, *dmCherry-P2A-CRE-, EYFP+*) (**Fig. 2E, F Supplementary Fig. S3C**). This classification encompassed 99.9% of mesenchymal cells. We first observed that *Shox2 initiating* cells were primarily found in progenitor clusters (**LP** and **EPP**) (**Fig. 2E, F**). This finding is also supported by a higher ratio of immature *Shox2* transcripts as measured by velocity analysis (La Manno et al., 2018) (**Fig. 2G**). Cells in the *maintaining* phase were located in proximal limb clusters, with a significant number also expressing *dmCherry-P2A-CRE* in addition to *Shox2* and *EYFP* (**Fig. 2E**). *Decommissioned* cells were in both proximal and distal clusters supporting the whole transcriptome velocity analysis (**Fig. 2C**) and the existence of distal limb cells originating from *Shox2*-expressing limb progenitors. Finally, *inactive* cells were predominantly observed in distal clusters (**Fig. 2E, F**). Over time, the proportion of *initiating* cells decreased concomitantly with an increase of *maintaining* and *decommissioned* cells (**Supplementary Fig. S3C**). In summary, while proximal clusters maintain *Shox2* expression initiated in limb progenitors, distal clusters increasingly decommission the locus transcription.

### The dual-color reporter approach allows sorting of *inactive*, *maintaining* and *decommissioned* cells over time

To further characterize the dynamics of the *Shox2* regulatory landscape, we utilized Fluorescence-Activated Cell Sorting (FACS) on *Shox2^trac^*fore- and hindlimb cells from E10.5 to E14.5. Yet, before conducting chromatin analyses, we evaluated through bulk RNA-seq, performed on the different FACS-sorted populations (**Fig. 3A**), whether sorted cells matched the *Shox2* transcriptional phases previously identified in the single-cell approach (**Supplementary Table S3**). We identified three major distinct cell populations. First, cells lacking fluorescence marker activity, classified as *inactive*, displayed no *Shox2* expression at E11.5 but expressed non-mesenchymal (*Krt5*, *Wnt6*, *Chd5*, *Myod1*) and mesenchymal *(Twis1, Msx1)* markers with a distal identity *(Hoxd13)* (**Fig. 3A, B, Supplementary Table S3**). Second, cells marked by high fluorescent levels of both dmCherry and EYFP, classified as *maintaining*, expressed high levels of *Shox2* and displayed a mesenchymal identity signal already at E11.5 (*Prrx1*, *Twist1*). Third, cells with high fluorescent levels of EYFP but no dmCherry signal, classified as *decommissioned*, were found not to express *Shox2* and to bear a mesenchymal identity (**Fig. 3A, B, Supplementary Table S3**). Additionally, we observed three minor cell populations. One was composed of dmCherry-only cells that strongly expressed blood markers genes such as *Hbb-y* and were therefore considered as being mainly blood cells. A second population comprised with low levels of both dmCherry and EYFP, termed “low-EYFP”, exhibited weak *Shox2* expression and displayed a mesenchymal progenitor identity by expressing high level of *Msx1* (**Fig. 3A, B**) (Markman et al., 2023). Interestingly, low-EYFP cells were notably abundant at early stages as they accounted for 7.5% of cells in E10.5 hindlimbs, but their prevalence decreased to 1% by E12.5-E13.5 (**Fig. 3C**). Finally, we identified a group of cells with high levels of EYFP and intermediate levels of dmCherry, expressing *Shox2*, that we termed “intermediate” and that bear a mesenchymal identity (**Fig. 3A, B**). Among these populations, we were unable to clearly identify a distinct group of *Shox2 initiating* cells. This observation led us to hypothesize that such cells might be dispersed among the identified cell populations (low-EYFP, inactive), possibly due to the delayed translation of dmCherry and EYFP. In summary, using FACS we could clearly isolate *inactive*, *maintaining* and *decommissioned* cell populations, yet we could not identify a pure *initiating* cell population.

**Figure 3:**
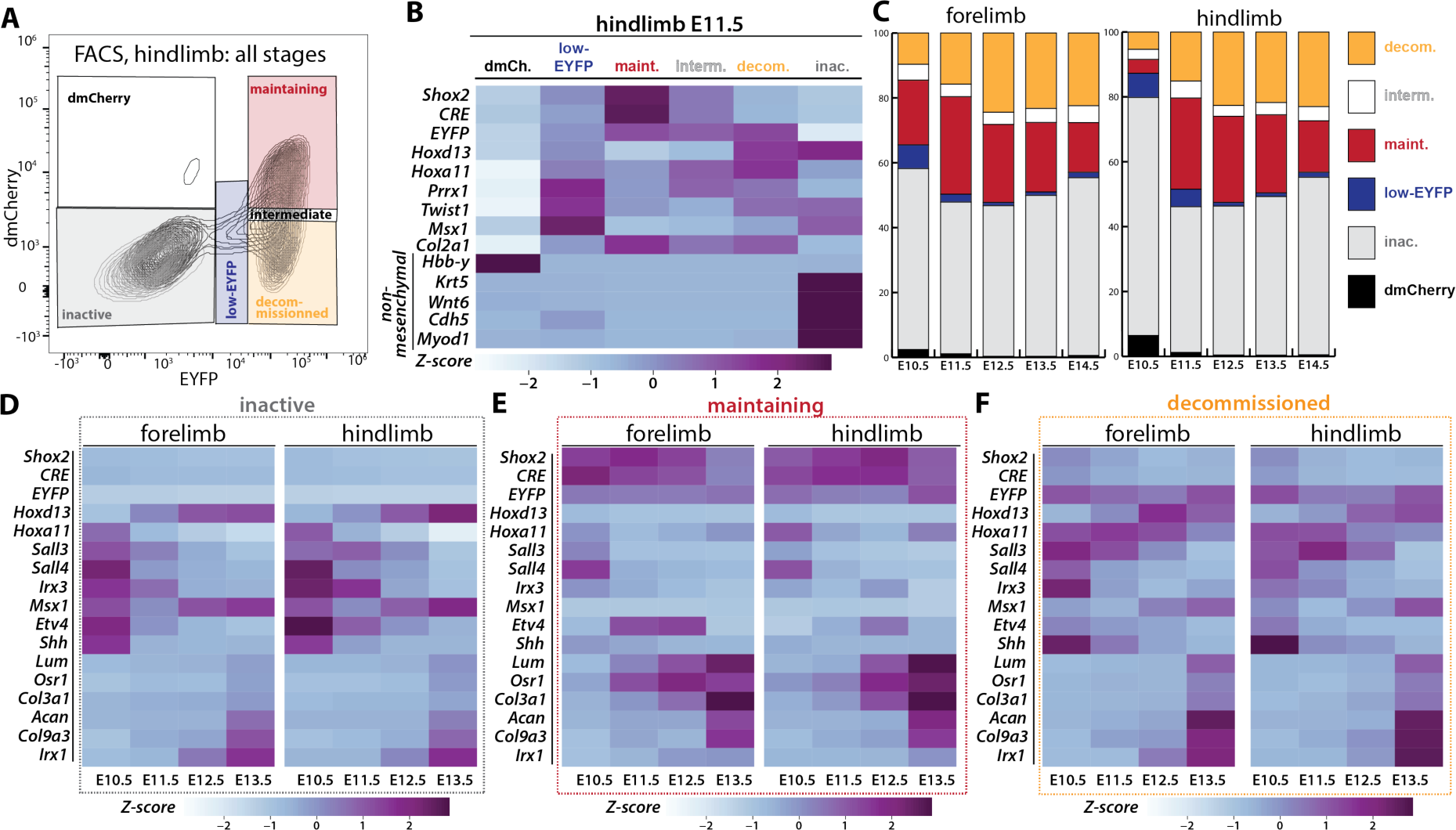
Characterizing the dual-color reporter approach to sort cells undergoing distinct phases of Shox2 regulatory trajectory. **A.** FACS plot of Shox2^trac^ hindlimbs dissociated cells illustrating the distribution of cells based on dmCherry and EYFP fluorescence in all the developmental stages investigated. **B.** Heatmap profiles of representative marker genes in different sorted populations from E11.5 hindlimbs. dmCh. = dmCherry, maint. = maintaining, interm. = intermediate and inac. = inactive. **C.** Proportions of each sorted population at E10.5, E11.5, E12.5, E13.5, and E14.5 in both forelimbs and hindlimbs, showcasing the dynamic changes in population sizes over developmental time. **D-F.** Heatmaps of gene expression analysis of marker genes in sorted cells throughout fore- and hindlimb development. **D.** Marker gene expression in inactive cells. **E.** Marker gene expression in maintaining cells. **F.** Marker gene expression in decommissioned cells. Inactive (EYFP-, dmCherry-, in grey), maintaining (EYFP+, dmCherry+, in red), decommissioned (EYFP+, dmCherry-, in yellow), blood (EYFP-, dmCherry+, in black), low-EYFP (low EYFP+, low dmCherry+, in blue), intermediate (high EYFP+, intermediate dmCherry+, in light grey or white). In **C**, **D-F**: Z-score scale derived from normalized FPKMs provides a normalized measure by rows of gene expression levels enabling comparison across samples.

As *inactive*, *maintaining* and *decommissioned* cell populations accounted for 83-95% of the cells at the stages analyzed, we focused on these three populations for further analyses (**Fig. 3C**). These were also the main populations identified in our single-cell analysis together with the initiating population (**Fig. 2E, F**). *Inactive* cells (EYFP-, dmCherry-) constituted 56% and 73% of fore- and hindlimb cells at E10.5, respectively, and first displayed a proximal (*Hoxa11*+, *Hoxd13*-) and progenitor (*Irx3*+, *Msx1*+) identity (**Fig. 3C, D**, **Supplementary Table S3**). At later stages, the proportion of *inactive* cells decreased to around 50% while shifting to a more distal identity (*Hoxa11*-, *Hoxd13*+) along with a gradual increased expression of chondrogenic and digit markers (*Col9a2*+, *Irx1+*) (**Fig. 3C, D**, **Supplementary Table S3**). The difference between fore- and hindlimbs proportion at early stages can be attributed to the developmental advance of forelimbs, where more cells have already initiated *Shox2* transcription by E10.5. In fore- and hindlimbs, *maintaining* cells (EYFP+, dmCherry+) represented 20% and 4% at E10.5, respectively, then increased up to 25-30% at E11.5-E12.5 while ultimately decreasing to 15% at later stages (**Fig. 3C**). This also underlines the developmental advance of forelimbs in terms of *Shox2* activation. *Maintaining* cells continuously expressed proximal markers (*Hoxa11*+) and rapidly differentiated into chondrogenic (*Col9a3+, Acan+)* and connective tissue (*Osr1+, Lum+, Col3a1+)* lineages (**Fig. 3E**, **Supplementary Table S3**). *Decommissioned* cells (EYFP+, dmCherry-) were rare at early stages, constituting 10% and 5% in E10.5 fore- and hindlimbs, respectively, and displayed a proximal (*Hoxa11*+, *Hoxd13*-) and progenitor (*Irx3*+, *Shh+, Etv4+*) identity. At later stages *decommissioned* cells progressively differentiate into connective tissue and cartilage (*Lum*+, *Col9a2*+) in both distal and proximal limb segments (*Hoxa11*+, *Hoxd13*+, *Irx1+*) (**Fig. 3F, Supplementary Table S3**). Generally, the changes in marker genes expressed in *inactive*, *maintaining* and *decommissioned* cell populations over time mirrored scRNA-seq and microscopy analyses (See **Fig. 1** and **2**).

Changes in the proportions of each cell phase over time (**Fig. 3C**) indicate that during early stages the activation of the *Shox2* locus reduces the pool of *inactive* cells to expand the population of *maintaining* cells which ultimately lead to an increase of *decommissioned* cells. Discrepancies in cell proportion identified by FACS, compared to single-cell analysis (see **Supplementary Fig. S3C**), can be explained by the inherent differences in sensitivity between the two techniques. In conclusion, the dual-color approach enables us to efficiently track and sort *inactive*, *maintaining*, and *decommissioned* cells across time and through differentiation processes. Furthermore, our findings suggest an early wave of *Shox2* decommissioning in *Irx3*+ progenitor cells, likely associated with the emergence of a distal limb progenitor population (*Msx1+*) that gives rise to anterior digits (*Shh+*). These results pave the way for defining how *Shox2* expression is maintained over time and through differentiation and ultimately decommissioned in cells that will compose the future distal limb.

### *Shox2* transcriptional maintenance is associated with distinct enhancer repertoires acting over time

Both single-cell analyses and RNA-seq data from FACS experiments have demonstrated dynamic changes in the limb cell types in which *Shox2* is transcriptionally active, progressing from limb progenitors to chondrogenic and connective cell types (see **Figs. 2, 3**). To support its transcription through these transitions and adapt to this changing environment, the *Shox2* regulatory landscape likely employs distinct sets of enhancers (see **Fig. 1**). To delineate the stage-specific enhancer repertoires of *Shox2*, we leveraged our reporter system to focus on the chromatin status of *Shox2 maintaining* cells. We generated H3K27ac ChIP-seq profiles, indicative of active enhancers, in E10.5, E11.5, E12.5, E13.5 forelimbs *maintaining* FACS sorted cells (dmCherry+, EYFP+) in (**Fig. 4A**)(Rada-Iglesias et al., 2011). The usage of *Shox2* expressing cells across four stages allowed to produce a highly accurate map of *Shox2* putative enhancers and to define time-windows of activity. Here, we identified 34 H3K27ac-marked enhancers within the *Shox2* TAD (mm39, chr3:66190000-67290000). While a majority of these enhancers displayed an activity restricted to early stages of limb development (early enhancers: 26/34) other showed consistent H3K27ac coverage across all stages (common enhancers: 4/34) or restricted to late stages (late enhancers: 4/34) (**Fig. 4A, Supplemental Table S4**). H3K27ac ChIP-seq profiles form E11.5, E12.5 and E13.5 hindlimbs *maintaining* cells showed an identical distribution of enhancer and a highly similar temporal restriction of their activities (**Supplementary Fig. S4**). Prior studies have tested 13 of these enhancers *in vivo*, with five driving limb reporter activity at specific embryonic stages, validating our approach for enhancer identification (**Fig. 4B, Supplementary Table S4**) (Abassah-Oppong et al., 2023; Osterwalder et al., 2018). These findings suggest that a significant number of early enhancers is required to initiate and maintain *Shox2* transcription during the early stages of limb budding, while fewer are necessary to sustain expression at later stages.

**Figure 4:**
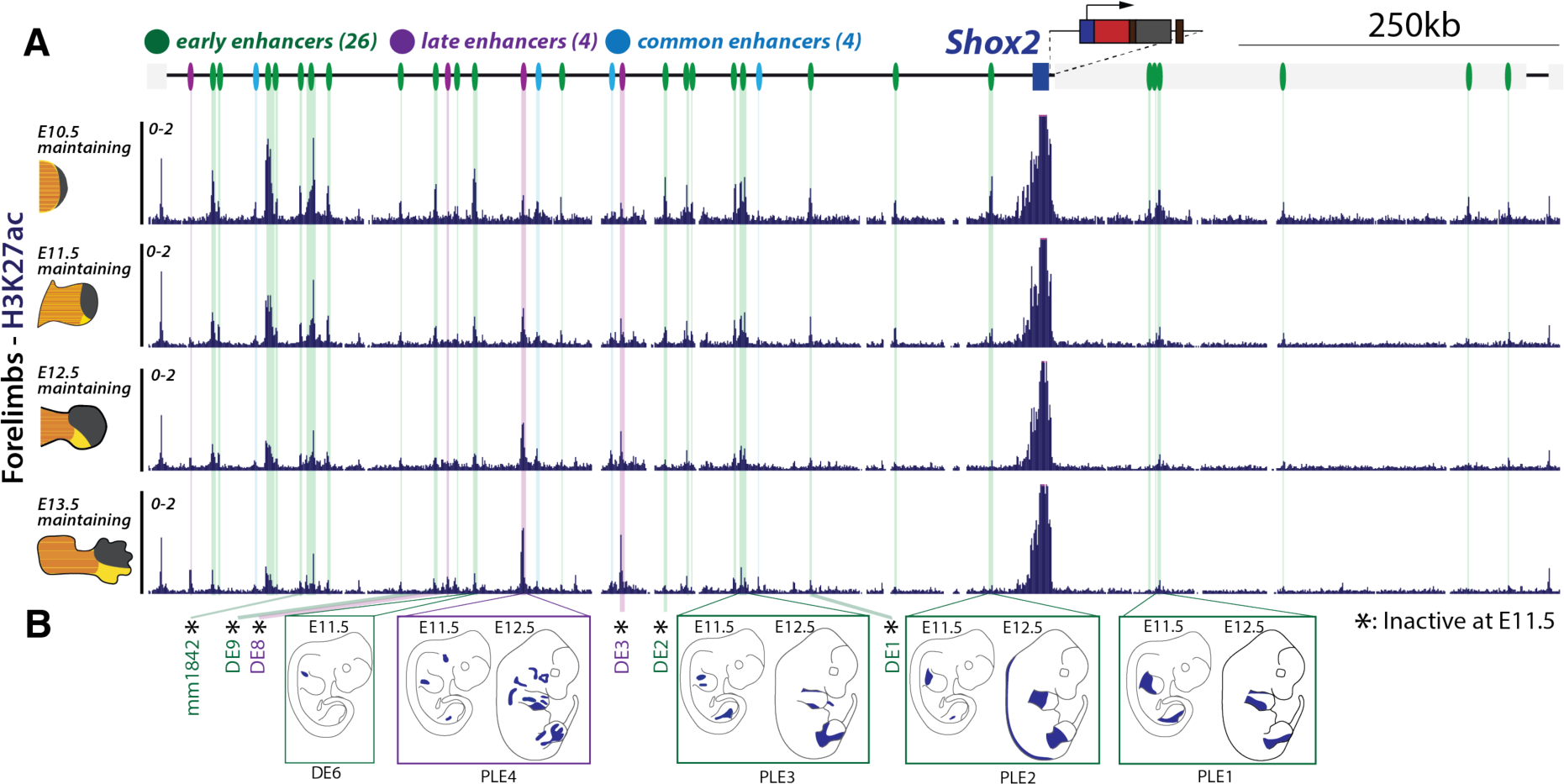
Regulatory maintenance associates with changes of enhancer repertoires. **A.** H3K27ac ChIP-seq profiles at the Shox2 locus (mm39: chr3:66,190,000-67,290,000) of FACS sorted maintaining (dmCherry+/EYFP+) cells across E10.5, E11.5, E12.5, and E13.5 forelimbs. Putative enhancers are shown by color-coded lines: green for early, light blue for common, and purple for late enhancers, as detailed **in Supplementary Table S4**. The light gray box next to the Shox2 gene is the Rsrc1 gene. **B.** Schematic representation of the pattern displayed by enhancers previously validated through in vivo LacZ reporter assays (Abassah-Oppong et al., 2023; Osterwalder et al., 2018).

### The 3D locus topology of *Shox2* mirrors temporal enhancer repertoire shifts

Recent studies have demonstrated that changes in chromatin architecture associates with the activity of enhancers and promoters (Robson et al., 2019; Rouco et al., 2021). We sought to investigate whether the temporal shifts in enhancer activities observed at the *Shox2* locus are also associated with temporal changes in the 3D chromatin organization. To tackle this question, we generated capture-HiC (C-HiC) maps across different phases and stages of the *Shox2* regulatory trajectory. We started by comparing *Shox2^trac^* embryonic stem cells (mESCs) with *inactive* (dmCherry-, EYFP-) fore- and hindlimb FACS sorted cells at E11.5. Notably, while the poised *Shox2* locus in mESCs exhibited a relatively relaxed structure, with few focal interaction contacts, increased contacts between the TAD boundaries and between *Shox2* and three of its early enhancers were observed in *inactive* cells (**Fig. 5A-C, Supplementary Fig. S5**) (Mas et al., 2018). We then studied transition from a poised state towards an active status, by comparing *Shox2^trac^* mESCs and fore- and hindlimb FACS-sorted *Shox2 maintaining* cells at E11.5, E12.5, and E13.5. We observed that in *maintaining* cells, unlike in *inactive* ones, interactions between *Shox2* and most of its enhancers were heightened (**Fig. 5A, C-E, C’ and Supplementary Fig. S5**). This was accompanied by a pronounced segregation of the locus into two subTADs in-between the *Shox2* and *Rsrc1* promoters. Remarkably, at E11.5, we noted increased interactions primarily with early and common putative enhancers, but not with late ones (**Fig. 5A, C, C’ and Supplementary Fig. S5**). This observation became more noticeable when comparing the locus structure in *maintaining* cells at E11.5 and E13.5 (**Fig. 5E, 5E’**). Specifically, this comparison revealed subtle yet consistent changes in contact patterns, where two early enhancers showed decreased interactions with *Shox2* at E13.5, while one and two late enhancers in fore- and hindlimbs, respectively, showed increased interactions (**Fig. 5A, C, Supplementary Fig. S5**). Thus, these observations show that the 3D structure of the *Shox2* locus accompanies a shift between early and late enhancer repertoires.

**Figure 5:**
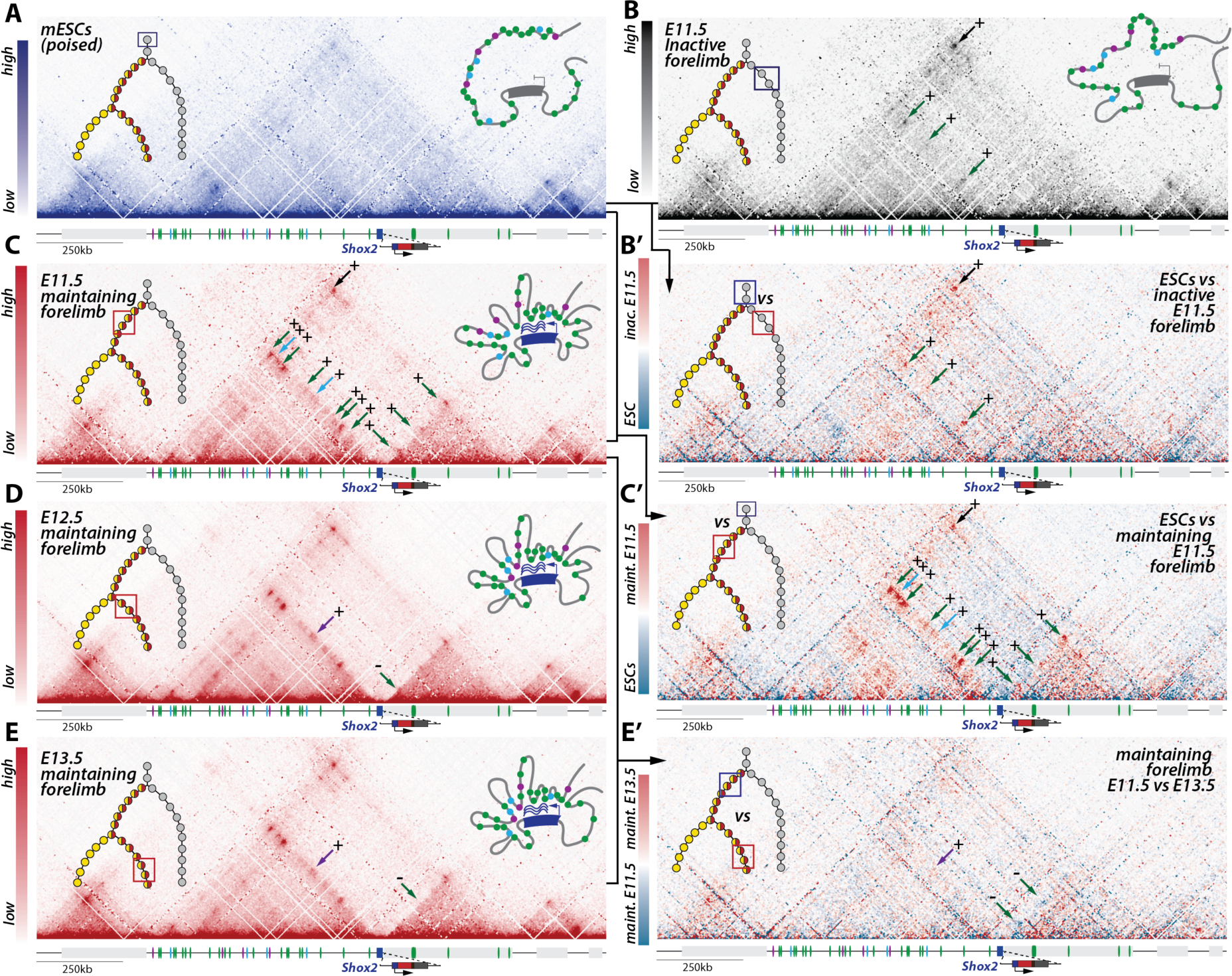
Shox2 locus 3D topology associates with active enhancer-promoter interactions in forelimbs. In all Capture-HiC (C-HiC) maps (mm39: chr3:65,885,132-67,539,263), the upper left illustration represents the position of the investigated cells in the regulatory trajectory and the upper right one a model of the average 3D locus structure. The light grey box next to Shox2 is the Rsrc1 gene. **A.** C-HiC maps of the Shox2 locus in Shox2^trac^ mESCs. Note a large TAD with few focal interaction points. **B.** C-HiC maps of the Shox2 locus in E11.5 forelimb FACS-sorted inactive cells. Note the formation of specific contacts with three early enhancers (green arrows) and increased loop contact between the two TAD borders (black arrow). **B’**. Subtraction C-HiC map between Shox2^trac^ E11.5 forelimb FACS-sorted inactive cells and Shox2^trac^ mESC. **C-E.** C-HiC maps of the Shox2 **C.** E11.5 **D.** E12.5, and **E.** E13.5 forelimb FACS-sorted maintaining cells. **C’**. C-HiC subtraction maps between Shox2^trac^ mESCs and Shox2^trac^ E11.5 forelimb FACS-sorted maintaining cells. **E’** C-HiC subtraction maps between E12.5 and E13.5 Shox2^trac^ FACS sorted forelimb maintaining cells. Changes in enhancer-Shox2 interactions are marked by colored arrows at each stage: green for early enhancers, purple for late enhancers, and light blue for common enhancers. A plus sign (+) denotes a gain of interaction, and a minus sign (-) indicates a loss of interaction relative to the previous stage. Also note the increased separation between the two subTADs at the position of the Shox2 gene body. Maps coordinates mm9; chr3:65,781,633-67,435,852. Maint. = maintaining; inac. = inactive.

### Shox2 enhancers act in an early and late specific manner

To functionally assess the role, timing and interdependency of identified early and late enhancers, we investigated whether deletions of these enhancers lead to transient, stage-specific alterations in *Shox2* expression. We first assessed the role of several early enhancers by generating two consecutive deletions, one of 139kb and another of 307kb, near the *Shox2* locus. These deletions removed 8 out of the 26 early enhancers (31%) in the *Shox2^trac^*background, while leaving the common and late enhancers intact (**Fig. 6A**). This double deletion allele, referred to as *Shox2^Δearly^ (Shox2^dmCherry/+;Δearly/Δearly^*; *Rosa26^loxEYFP/+^)*, was examined using fluorescent imagining, quantification of the proportion of *inactive*, *maintaining* and *decommissioned* cells as well as RNA-seq. At E11.5

**Figure 6:**
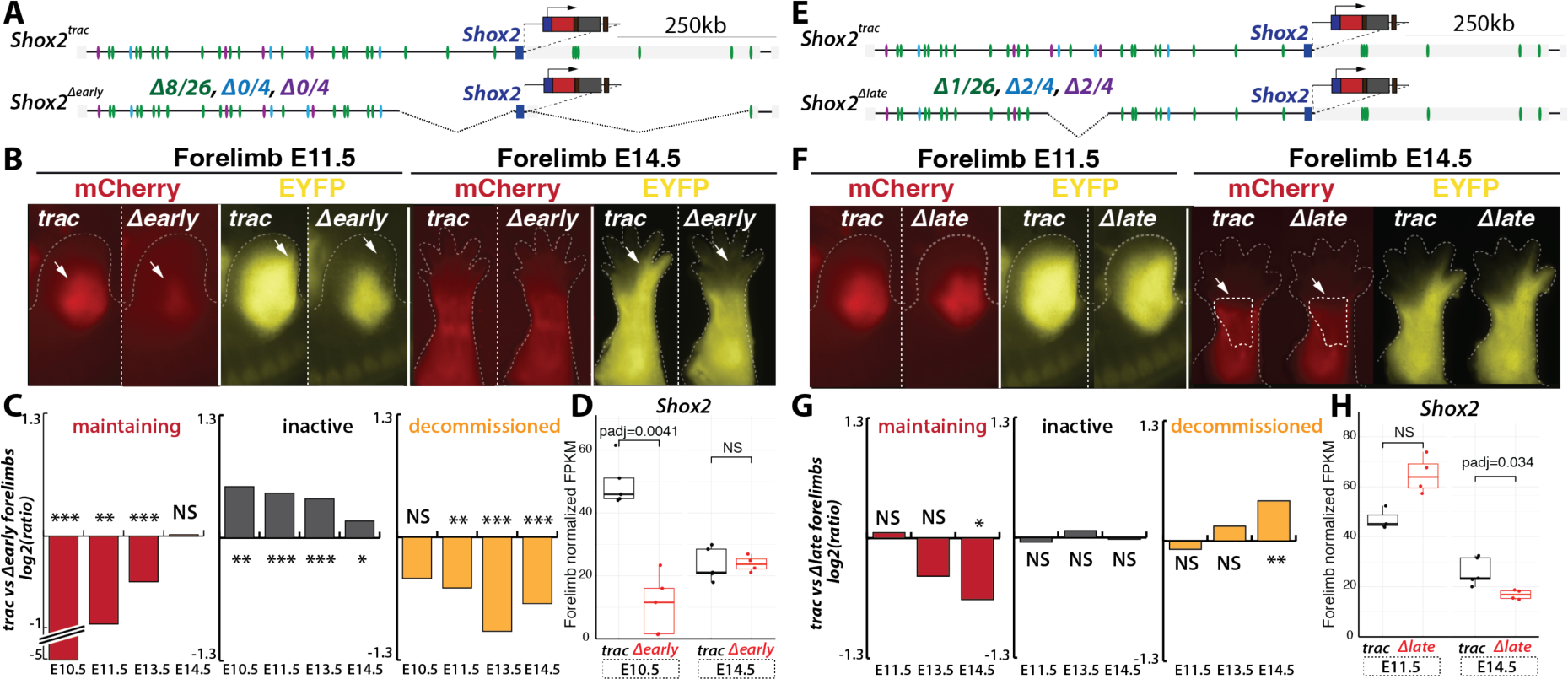
Partial early and late enhancer deletion induce stage-specific alterations. **A.** Illustration detailing the Shox2^Δearly^ deletion allele lacking 8 out of 26 early putative enhancers (in green), while late (in purple) and common (light blue) putative enhancers remain intact. **B.** Fluorescent imaging comparing Shox2^trac^ and Shox2^Δearly^ forelimbs at E11.5 and E14.5. White arrows indicate a reduction in dmCherry and EYFP signals at E11.5 and a complete loss of EYFP signal in distal limbs at E14.5. **C.** Log2 ratio of the proportion of maintaining, inactive, and decommissioned cell populations, identified by flow cytometry analyses, in Shox2^trac^ versus Shox2^Δearly^ forelimbs at E10.5, E11.5, E13.5, and E14.5. **D.** Normalized FPKM of Shox2 expression in E10.5 and E14.5 Shox2^trac^ and Shox2^Δearly^ entire forelimbs, with comprehensive DESeq2 analyses in **Supplementary Table S6**. **E.** Schematic representation of the Shox2^Δlate^ deletion allele lacking 2 out 4 late (in purple), 2 out of 4 common (in light blue), 1 out of 26 early putative enhancers. **F.** Fluorescent imaging of Shox2^trac^ and Shox2^Δlate^ forelimbs at E11.5 and E14.5. White arrows and delimited area indicate a loss of dmCherry signal in the central limb section at E14.5. **G.** Log2 ratio of the proportion of maintaining, inactive, and decommissioned cell populations, identified by flow cytometry analyses, Shox2^trac^ versus Shox2^Δlate^ forelimbs at E11.5, E13.5, and E14.5. **H.** Normalized FPKM of Shox2 in E10.5 and E14.5 Shox2^trac^ and Shox2^Δlate^ entire forelimbs, detailed DESeq2 analyses in **Supplementary Table S6.** In **C** and **G**: T-tests were utilized to calculate p-values from replicates (**Supplementary Fig. S6**). NS= non-significant, *=p<0.05, **=p<0.01 and ***=p<0.001. In **D** and **H** boxplots, boxes indicate the first and third quartiles, the whiskers indicate ±1.5 × interquartile range, and the horizontal line within the boxes indicates the median. Statistical test used: DESeq2 Wald test, padj is the FDR-corrected p-value from DESeq2.

*Shox2^Δearly^* embryos displayed a limb-wide decreased in dmCherry and EYFP signals compared to those in *Shox2^trac^* forelimbs (**Fig. 6B**). We also observed the loss of the distal limb EYFP signal, not overlapping dmCherry, in *Shox2^Δearly^*limbs. By E14.5, dmCherry changes were no longer apparent, but the loss of EYFP signal in distal segments persisted. Quantification of cell proportions from flow cytometry experiments in *Shox2^Δearly^* fore- and hindlimbs at E10.5, E11.5, E13.5, and E14.5 showed a complete loss of *maintaining* cells at E10.5 (33 fold less, from 19% in *Shox2^trac^* forelimb cells to 0.6% in *Shox2^Δearly^*) that gradually returned to control levels at later stages (**Fig. 6C, Supplementary** Fig. S6A, S7A, B**, Supplementary Table S5**). Additionally, we observed an increase in *inactive* cells, especially at early stages (1.5 fold more: from 56% in *Shox2^trac^* forelimbs to 81% in *Shox2^Δearly^*), likely due to the delayed *Shox2* transcriptional onset. Concomitantly, we noted a decrease in *decommissioned* cells reflecting the observed loss of distal EYFP signal (for instance at E13.5 2 fold less: from 23.3% in *Shox2^trac^* forelimbs to 11.7% in *Shox2^Δearly^*, **Fig. 6B, C, Supplementary** Fig. S6A, S7A, B**, Supplementary Table S5**). RNA-seq of E10.5 and E14.5 forelimbs revealed a decrease in *Shox2* expression only at E10.5, aligning with the changes observed in dmCherry fluorescence (**Fig. 6D, Supplementary Table S6**). These findings indicate that the removal of early enhancers leads to a strong but transient decrease in *Shox2* transcription in early limb progenitor cells. While transcription in proximal lineages is gradually restored by the remaining enhancers, cells destined for distal posterior limb segments do not regain *Shox2* expression (see **Fig. 2**).

Conversely, we hypothesized that late enhancers, in conjunction with common ones, are crucial for sustaining *Shox2* expression during the late stages of limb development, without significantly contributing to the early phase. To test this hypothesis, we engineered a targeted deletion spanning 84kb, *Shox2^Δlate^ (Shox2^dmCherry/+;Δlate/Δlate^*; *Rosa26^loxEYFP/+^)*, that eliminates 2 out of 4 late enhancers (50% of late), 2 out of 4 common enhancers (50% of common) and 1 out of 26 early enhancers (4% of early) (**Fig. 6E**). At E11.5, no discernible differences in dmCherry or EYFP fluorescence were observed in forelimb tissues compared to controls (**Fig. 6F**). However, a slight decrease in dmCherry signal (but not EYFP) was noted in the central section of E14.5 forelimbs (**Fig. 6F**). Flow cytometry analysis of cell proportions in *Shox2^Δlate^*forelimbs at E11.5, E13.5, and E14.5 revealed a specific reduction in *maintaining* cells solely at E14.5 (1.6 fold less: from 15% in *Shox2^trac^* to 10% in *Shox2^Δlate^*, **Fig. 6G, Supplementary Fig. S6B**). RNA-seq analysis at E11.5 and E14.5 confirmed a loss of *Shox2* expression exclusively at E14.5 (**Fig. 6H, Supplementary Table S6**). Unlike the early enhancer deletion, the late enhancer deletion did not impact the proportion of *inactive* cells but only led to an increase in *decommissioned* cells (1.3 fold more: from 22% in *Shox2^trac^*to 30% in *Shox2^Δlate^*), indicating that previously ongoing transcription was halted due to the absence of late regulatory elements, entering in a premature decommissioning phase (**Fig. 6G**). Surprisingly, hindlimb analyses showed no significant changes, possibly due to the slight developmental delays between fore- and hindlimbs (**Supplementary Fig. S7C, D**). Together, these deletion experiments demonstrate that *Shox2* transcription is regulated by different enhancer repertoires operating transiently, in a temporal-specific manner during limb development. Moreover, it suggests that early and late enhancers might act in an independent way.

### Enhancer-Promoter disconnection marks *Shox2* decommissioning

Despite their fundamental contribution to the formation of gene expression patterns, the features associated with the termination of a locus’ transcriptional activity remain poorly characterized. To define how *Shox2* transcription is terminated, we examined the changes in locus topology, transcription, and activities of regulatory elements within *decommissioned* cells. First, we generated C-HiC maps of FACS-sorted *decommissioned* cells from *Shox2^trac^* E12.5 and E13.5 fore- and hindlimbs (**Fig. 7A, Supplementary** Fig. S8A, B, C). Compared to stage-matched *maintaining* cells, we observed a notable reduction in enhancer-promoter interactions and a decrease in the segregation of the two subTADs. We noticed that this decommissioned structure was very similar to the one seen in *inactive* cells (as shown in **Fig. 5B, Supplementary Fig. S8D**). Looking at the locus regulatory activity, we observed a significant decrease in *Shox2* expression and H3K27ac coverage of the gene’s promoter in FACS-sorted *decommissioned* cells compared to *maintaining* ones (**Fig. 7B, Supplementary Fig. S9A**). At the enhancer level, we noted a faster depletion of H3K27ac coverage than at the *Shox2* promoter (**Fig. 7C, Supplementary Fig. S9B**), especially visible at early and common enhancers. However, by the late E13.5 stage, two late enhancers (see **Fig. 4B**) still retained H3K27ac coverage in *decommissioned* cells. Intriguingly, these same regions also exhibited some activity in late E12.5 and E13.5 *inactive* cells (**Supplementary Fig. S9C**). This suggests that these two enhancers, although significantly contributing to *Shox2* maintenance (see **Fig.6 E-H**), are insufficient to sustain or initiate *Shox2* expression by themselves. Taken together, these findings indicate a rapid disconnection of the majority of enhancer-promoter contacts, accompanied by a swift reduction in their active chromatin coverage during locus decommissioning.

**Figure 7:**
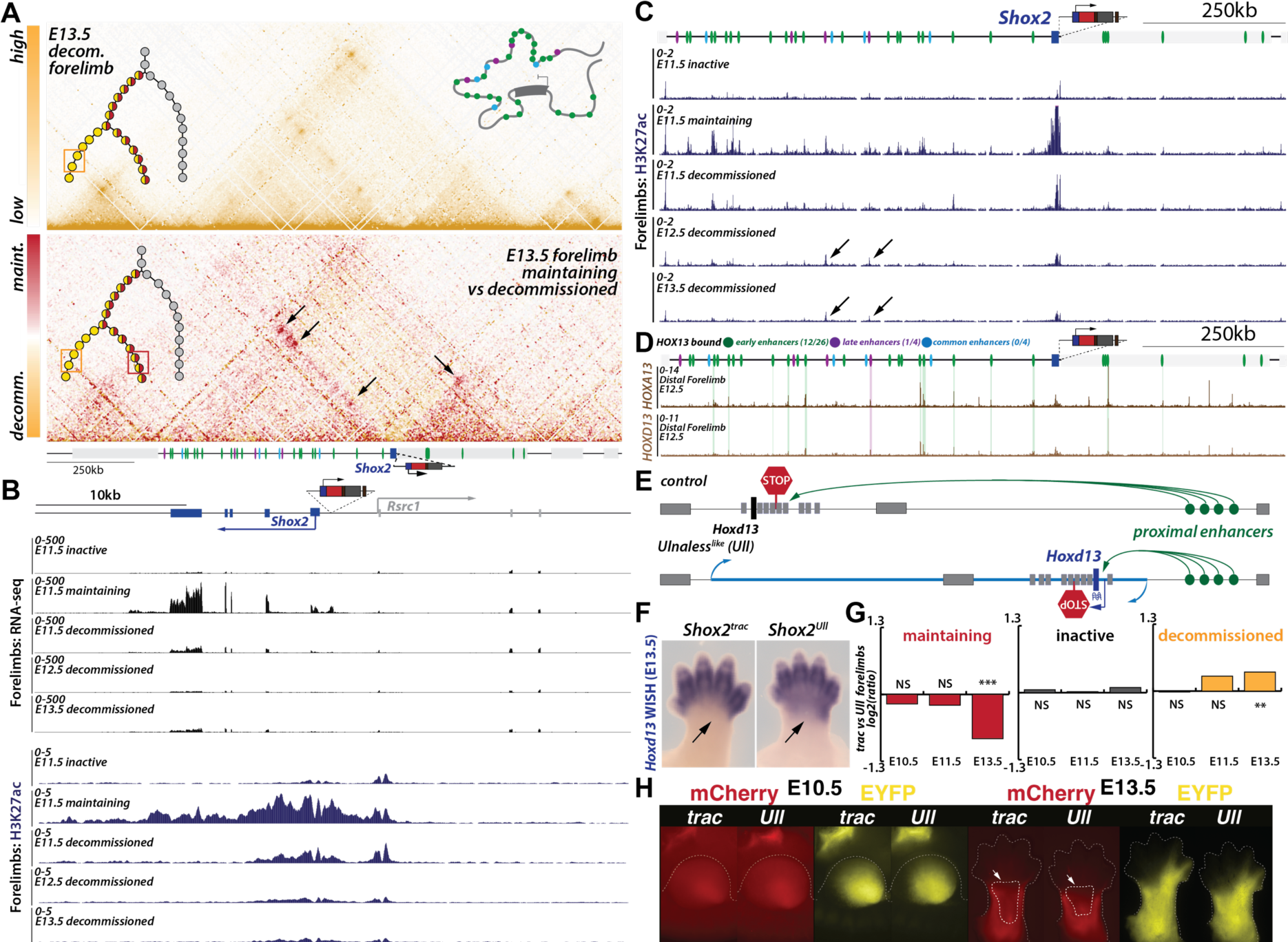
Shox2 locus decommissioning associate with the disconnection between enhancers and promoters. **A.** C-HiC of E13.5 forelimb FACS-sorted decommissioned cells (above) and subtraction map with E13.5 forelimb FACS-sorted maintaining cells (See Fig. 5C); in each panel the upper left illustration represents the position of the investigated cells in the regulatory trajectory; the upper right one is a model of the average 3D locus structure (maps coordinates mm39; chr3:65,885,132-67,539,263). Black arrows indicate strong losses of enhancer-promoter inteactions **B.** RNA-seq and H3K27ac ChIP-seq tracks of early (E11.5) forelimb inactive, maintaining, decommissioned and late (E12.5-E13.5) decommissioned FACS-sorted cells at the Shox2 and Rsrc1 promoter region (mm39: chr3:66,870,000-66,910,000). **C.** Forelimb H3K27ac ChIP-seq tracks in early (E11.5) forelimb inactive, maintaining, decommissioned and late (E12.5-E13.5) decommissioned FACS-sorted cells over the Shox2 regulatory landscape (mm39: chr3:66,190,000-67,290,000). Note the overall loss of H3K27ac at enhancers in decommissioned cells while two of the four late enhancers show activity in decommissioned cells (black arrows). **D.** HOXA/D13 ChIP-seq in distal forelimb at E12.5 re-analyzed from (Sheth et al., 2016). **E.** Schematic illustration of the engineered Ulnaless inversion allele at the Hoxd locus, showing how the inversion brings proximal enhancers into close proximity with Hoxd13, leading to its misexpression.**F**. Hoxd13 WISH in wildtype and Shox2^Uln^ forelimbs at E13.5. Note how the Ulnaless inversion allele induce proximal expression of Hoxd13 in forelimbs (black arrow). **G.** Log2 ratio between the proportion of forelimb Shox2^trac^ and Shox2^Uln^ initiating, maintaining and decommissioned cell population, identified by flow cytometry at E10.5, E11.5 and E13.5. T-tests were utilized to calculate p-values from replicates (**See supplementary Fig. S10**). NS= non-significant, *=p<0.05, **=p<0.01 and ***=p<0.001. **H.** Fluorescent imaging of Shox2^trac^ (trac) and Shox2^Uln^ (Uln) forelimbs at E10.5 and E13.5. Note the decreased dmCherry signal in the central part of the limb (white arrow). Maint. = maintaining; decomm. = decommissioned.

### An *Ulnaless* like allele triggers ectopic *Shox2* locus decommissioning

The rapid disconnection of enhancer-promoter contacts and the swift decline in enhancer activities upon *Shox2* locus decommissioning imply a targeted repressive activity at enhancer regions, potentially mediated by the binding of specific transcription factors (TFs). Furthermore, the presence of many *decommissioned* cells in the distal limb domain suggests the involvement of distal limb TFs in this process (**Fig. 2, 3**). *HOXA/D13* TFs have been involved in *Shox2* repression in distal limbs and both factors are binding several identified *Shox2* enhancers (**Fig. 7D**) (Beccari et al., 2016; Sheth et al., 2016). Notably, out of 34 *Shox2* putative limb enhancers, *HOXA/D13* proteins were found to bind to 13 of them, encompassing 46% (12/26) of early acting enhancers and 1 out of 4 late enhancers in distal forelimbs at E12.5. This positions *HOXA/D13* as candidate TFs for controlling *Shox2* decommissioning by inhibiting its early expression phase in limb progenitors.

To explore this hypothesis, we employed a limb-specific gain-of-function allele known as *Ulnaless*, which induces *Hoxd13* expression in proximal limbs where *Shox2* is normally active (Bolt et al., 2021; Herault et al., 1997). Here, we re-engineered the *Ulnaless* inversion (*Ulnaless-like (Ull)*) in the dual-color tracking background, *Shox2^Ull^* (*Shox2^dmCherry/+;^*; *Rosa26^loxEYFP/+^*; *HoxD^Ull/+^*) (**Fig. 7E**). As expected, *Shox2^Ull^* displayed a gain of *Hoxd13* transcripts in the proximal region of E13.5 forelimbs (**Fig. 7F**). If *Hoxd13* acts as a decommissioning factor, we would expect to observe a decrease in *maintaining* cells and an increase in *decommissioned* cells, without affecting *inactive* cells. Quantification of such cell proportion of cells by flow cytometry in both fore- and hindlimbs precisely revealed a decrease in *maintaining* cells (1.8 fold less, from 21% of forelimb cells in *Shox2^trac^* to 12% in *Shox2^Ull^*) and an increase in *decommissioned* cells (1.3 fold more, from 23% in *Shox2^trac^* forelimb to 30% in *Shox2^Ull^*) at the late E13.5 stage, but not at earlier stages (**Fig. 7G, Supplemental Fig. S9D, S10)**. Additionally, fluorescent microscopy revealed a mild reduction of dmCherry signal in the center of E13.5 forelimbs but not at E11.5 (**Fig. 7H**). These findings mirror the effects observed by late enhancer loss (See **Fig. 6**) and demonstrate that *HOXD13* can induce locus decommissioning, likely through binding active enhancers.

## Discussion

Developmental genes exert their influence through spatially and temporally restricted expression patterns (Cao et al., 2019; Chen et al., 2022; He et al., 2020). Building upon previous studies on enhancer dynamics, which explored enhancer activation and decommissioning, we delve into the behavior of an entire gene regulatory landscape over time (Maeda and Karch, 2006; Whyte et al., 2012; Wu et al., 2023). We conceptualized the study of this regulatory dynamics, affecting gene expression across extended periods as a regulatory trajectory (see **Fig. 1D**). Through our investigation, with *Shox2* serving as a testbed locus, we reveal that in the early stages of limb development, cells either remain inactive, contributing predominantly to distal limb cell populations, or become transcriptionally active initially in the limb progenitors. This activity is subsequently maintained during differentiation into proximal connective tissue and cartilage. After transcriptional initiation and maintenance, we identified that certain cells, including some destined to become part of distal limb segments, undergo decommissioning of the *Shox2* locus, possibly through the activities of distally expressed TFs such as HOXA/D13.

During early limb development, we have observed that limb progenitors seem to be neither proximal nor distal as they express both *Hoxd13* and *Shox2*. Previous studies have also remarked the same observation, even suggesting that these cells could contribute to the mesopodium (wrist-ankle) formation (Desanlis et al., 2020; Markman et al., 2023; Neufeld et al., 2014). Using our tracing system, we were able to precisely track the fate of cells that express *Shox2* in early stages but cease to do so later. We found that *decommissioned* cells not only contribute to the mesopodium but also to proximal and distal limb cell populations, including digits. Early *decommissioned* cells exhibited expression of the *Msx1* limb progenitor marker, along with *Irx3/5* and *Shh* genes, which are crucial for digit development (Li et al., 2014; Zhu et al., 2022). Moreover, the removal of one-third of the early enhancers led to a delayed *Shox2* expression initiation and, subsequently, a loss of *Shox2 decommissioned* cells in distal limbs. These outcomes show that *Shox2* expressing limb progenitors that rely on the activities of early enhancers contribute to the formation of the autopod and in particular of digits 4 and 5.

Curiously, despite the significant delay and loss of *Shox2* expression following early enhancer deletion, the late expression eventually manifests, indicating the capacity of remaining early, common, and late enhancers to sustain proximal *Shox2* expression at late stages. Yet, it remains uncertain whether the restoration of expression levels results from a collective effect of all enhancer classes or is attributable to specific ones. Additionally, whether late enhancers alone can restore expression following the complete abrogation of early enhancers - and what the resulting expression pattern might look like - has yet to be clarified. Insights from the late enhancer deletion experiment, which led to an increase in *decommissioned* cells without affecting inactive cells, specifically at late stages, suggest late enhancers can only act on a previously activated *Shox2* locus and cannot activate expression in inactive cells. This hypothesis is further supported by the presence of H3K27ac on some late enhancers in *inactive*, *Shox2*-cells. Hence, these enhancers likely influence the locus only in cells already imprinted with the patterning cues provided by early enhancers.

Initially, during the onset of *Shox2* transcription, enhancer-promoter contacts are strongly increased alongside the formation of a subTAD boarder at the *Shox2* promoter position. As development progress, contacts appear to shift from early-to late enhancers, as the first are becoming inactive and the latter active. Finally, during decommissioning, enhancers are disconnected from *Shox2* simultaneously with the decrease in H3K27ac coverage. This highly dynamic structure cannot be explained by differential binding of CTCF, as the architectural protein does not bind the *Shox2* promoter at any stage and only displays minimal binding differences within the *Shox2* centromeric regulatory landscape (**Supplementary Fig. S11**)(Andrey et al., 2017). Here, two different CTCF-independent mechanisms might be at play to explain this dynamic enhancer-promoter contact behavior. First, it is conceivable, as recently proposed, that cohesin is loaded at enhancers via RNAPII and subsequently extrudes the local domain until hitting the *Shox2* promoter region. In turn, this would lead to a loading of cohesin at the promoter and a self-forward acting structural loop (Zhang et al., 2023). In this view, enhancer promoter contacts would be stabilized independently of CTCF but with a high dependency on enhancer activities. Alternatively, the contact might be depending on the recruitment of looping factors at the various active enhancers enabling the formation of micro-compartments independently of loop extrusion as recently shown in another study (Goel et al., 2023). After transcriptionally active phases, gene expression can rapidly shut down to establish sharp expression boarders critical for defining anatomical transitions. In the case of *Shox2*, its widespread expression in the early limb bud becomes restricted to proximal limb segments, with a sharp expression boundary at the mesopodium level (Neufeld et al., 2014). This boundary precisely opposes the expression onset of *Hoxa/d13,* marking the region where digits form. Here, we have shown how *Ulnaless-like* embryos present a premature decommissioning of the *Shox2* locus likely causing the alteration of *Shox2* boundaries of expression (Gross et al., 2012). This correlate with the proximal shift of the *Hoxd13* expression domain (see **Fig.7F**), and with the known role of HOXD/A13 in repressing the early/proximal phase of *Hoxd* genes themselves (Andrey et al., 2013; Andrey et al., 2017; Beccari et al., 2016). However, the exact mechanisms of this decommissioning remain elusive. We hypothesize that similar mechanisms to those observed during the exit from pluripotency might be in play, where the histone lysine-specific demethylase 1 (LSD1) and histone de-acetylases (HDACs) target and dismantle a significant fraction of pluripotency enhancers (Respuela et al., 2016; Whyte et al., 2012). This decommissioning involves the demethylation and deacetylation of H3K4me2 and H3K27ac, respectively, effectively reverting loci to a poised state that is permissive for either activation or terminal repression. Indeed, we observed that normal *Shox2* locus decommissioning is featured by a faster loss of active H3K27ac marks at enhancers than at the *Shox2* promoter. In this context, HOXD/A13 proteins could tether HDACs and/or LSD1 to all enhancers of proximal loci including *Shox2*. This deactivation could then trigger the disassembly of the locus 3D structure and particularly of enhancer-promoter contacts as observed in all *decommissioned* cells.

Overall, our findings underscore the existence of temporally restricted enhancer repertoires, their synchronized shutdown during locus decommissioning, and the significant structural transformations accompanying these regulatory events. These insights are potentially applicable to dozens of other limb-associated loci (see **Fig. 1**) and are likely extendable to genes involved in other morphogenetic processes.

### Limitation of the study

A challenge in this study was the inability to sort a distinct initiating population. This issue precluded a detailed analysis of the *Shox2* regulatory landscape during the transcriptional initiation phase. Therefore, further investigations are necessary to establish whether *Shox2* transcriptional initiation is influenced by all identified enhancers, only a subset, or other elements acting at an earlier stage.

## Material and Methods

### Animal procedures

Animal research was performed at University of Geneva following the institutional, state and government regulations (Canton de Genève authorizations GE/89/19 and GE192A).

### CRISPR/Cas9 engineered alleles

Alleles genetically engineered were edited using CRISPR/Cas9 and following a similar procedure as (Kraft et al., 2015). In brief, sgRNAs were designed using the online software Benchling (https://benchling.com/) and were chosen based on predicted on-target and off-target scores. All the sgRNAs used for this study, their CRISPR/Cas9 genomic target location and genotyping primers, can be found in **Supplementary Table S7**. Each sgRNA was cloned in the plasmid pX459 (Addgene, #48139) and 8µg of each vector was used during mESCs transfection following the standard procedure for mESCs culture and genomic editing (Andrey and Spielmann, 2017). To construct the *Shox2^dmCherry/+;RosaEYFP/+^* or *Shox2^trac^* mESC clone, two rounds of targeting on G4 male cells, obtained from the Nagy laboratory (http://research.lunenfeld.ca/nagy/?page=mouse%20ES%20cells), were performed, first to insert the dmCherry-CRE cassette at the *Shox2* locus (*Shox2^dmCherry^*^/+^),followed by a second round of targeting to insert the EYFP cassette at the *ROSA26* locus (*Shox2^trac^)*. To integrate the dmCherry-CRE cassette cells were transfected with 8µg of the corresponding sgRNA and 4µg of in house designed cassette containing unbalanced homology arms (1.7 kb and 0.5kb), minimal β-globin promoter, dmCherry reporter sequence, PEST sequence, P2A self-cleaving peptide, CRE protein sequence and a bGH-PolyA terminator. To integrate the EYFP cassette, the EYFP sensor from (Srinivas et al., 2001) was modified by shortening the homology arms (final length of unbalanced arms 1.4kb and 0.85kb) and by removing the PGK promoter, the Neo/Kan selection cassette, the polyA of the PGK and the first spacer before the first SV40 poly(A). In house designed and modified cassettes were synthesized by Azenta Genomics/GENEWIZ. Alleles containing deletions or inversions were created in subsequent rounds of targeting performed using the *Shox2^trac+^* mESC clone as starting clone.

### Aggregation of mESCs clones and embryo collection

Mouse embryos were obtained following the tetraploid complementation procedure (Artus and Hadjantonakis, 2011). In brief, two days before the aggregation procedure, desired clones were thawed, seeded on male and female CD1 feeders, and grown. Donor tetraploid embryos were provided from in vitro fertilization using c57bl6J x B6D2F1 backgrounds. Aggregated embryos were transferred into CD1 or B6CBA foster females. Animals were obtained from Janvier laboratories or from in house crosses. Embryos were collected in 1X DPBS (Gibco, 14190-094) at the desired stage depending on the downstream protocol. The presence of desired mutations in embryos was confirmed by PCR genotyping (Supplementary Table S7).

### Live fluorescent imaging

Embryos were imaged in 1X DPBS (Gibco, 14190-094) in a petri dish on a Zeiss Axio Zoom V16 using the Axiocam 506 color camera for brightfield images or the Axiocam 506 mono camera for fluorescent images, after fluorescent laser stimulation (Illuminator HXP 200C) using filter 46 HE YFP (excitation BP 500/25, emission BP 535/50) for capturing EYFP signal and filter 63 HE mRFP (excitation BP 572/25, emission BP 629/62) for capturing dmCherry signal. Images were taken using the Zen Blue Software v3.6. Adjustment of brightness was performed using Adobe Lightroom v6.4.

### Light sheet microscopy imaging

E12.5 embryo was fixed overnight in 4% PFA and storage in 1x PBS at 4 °C until clearing procedure started. The entire embryo was cleared using passive CLARITY based clearing method. Tissue was first incubated for three days at 4°C in a Bis-free Hydrogel X-CLARITY™ Hydrogel Solution Kit (C1310X, Logos Biosystems) to allow hydrogel solution diffusion into the tissue. This was followed by polymerization in a Logos Polymerization system (C20001, Logos Biosystem) at 37°C for 3 hours. The SDS-Clearing solution was prepared by dissolving 24.73g of boric acid (Sigma B7660 or Thermofisher B3750) and 80g of sodium dodecyl sulfate (Brunschwig 45900-0010, Acros 419530010, or Sigma L3771) in dH2O to make 2L of 4% SDS solution, adjusting the pH to 8.5. Samples were then washed twice for 30 minutes in PBS, immersed in the SDS-based clearing solution at 37°C for 48 hours, followed by two PBS washes with 0.1% TritonX. Finally, tissue was placed in a Histodenz© based-refractive index-matching solution (Histodenz Sigma D22158, PB + Tween + NaN3 pH 7.5 solution, 0.1% Tween-20, 0.01% NaN3, in 0.02 M phosphate buffer, final solution pH 7.5). Imaging was performed with a home-built mesoscale single-plane illumination microscope, the mesoSPIM microscope is described in (Voigt et al., 2019). In brief, the sample was excited with 488nm, 561nm and 647nm lasers. The beam waist was scanned using electrically tunable lenses (ETL, Optotune EL-16-40-5D-TC-L) synchronized with the rolling shutter of the sCMOS camera. This produced a uniform axial resolution across the field-of-view (FOV) of 5 μm. EYFP signal was filtered with 530/43 nm, dmCherry signal with 593/40 and far-red signal with LP663 bandpass filter (BrightLine HC, AHF). Z-stacks were acquired at 5 μm spacing with a zoom set at ×1.25 resulting in an in-plane pixel size of 5.26 μm. Background and autofluorescence signal were subtracted using the 561 nm excitation channel during images pre-processing. This step together with subsequent normalization and filtering of the images were performed with the Amira 2020.1 software. 3D videos and images were captured using the Imaris 9.6.0 software.

### Whole Mount In Situ Hybridisation (WISH)

#### Probes design and production

*Hoxd13* probe was produced by amplification from mouse wildtype DNA using primers located at the 3’UTR of the desired genes. Primers were designed with Prime3 Software v 0.4.0 (using default parameters except: product size range 400-600bp; primer size 15-19bp; primer Tm 60-62°), extended with either SP6 or T7 primer sequence (*Hoxd13* probe forward primer: CAAGCTATTTAGGTGACACTATAGTGCTGCCCAATCCGACT; *Hoxd13* probe reverse primer: GAACTGTAATACGACTCACTATAGGGCGTGCCTTCAACCTCCAA). *Shox2* probe was extracted from plasmid donated by John Cobb (Cobb and Duboule, 2005) by digestion with NcoI. After PCR amplification or plasmid digestion product was purified with Monarch PCR clean-up kit (NEB, #T1030s) and used to produce the DIG-labeled single-stranded RNA probes using the DIG RNA labeling kit (Roche, #11175025910). Probes were then cleaned with MegaClear Kit (Thermo Fisher, #AM1908).

#### WISH staining protocol

Embryos from the desired stage and genotype were fixed overnight in 4% PFA/PBS. Subsequently, the embryos were washed in PBST (PBS with 0.1% Tween), followed by dehydration in methanol/PBST solutions of increasing concentrations (30%, 50%, and 70%), with final storage at −20 °C in 100% methanol. For the WISH protocol, on the first day, embryos were bleached in 6% H_2_O_2_/PBST for 1 hour at room temperature (RT), followed by rehydration in reverse methanol/PBST steps, then washed in PBST. Embryos were then treated with 2µg/ml proteinase K/PBST for 3 minutes, incubated in 2mg/ml glycine/PBST, washed again in PBST, followed by three 30 minutes washes in RIPA buffer (5M NaCl; 10% NP-40; 10% Deoxycholate; 20% SDS; 500mM EDTA pH8; 1M Tris-HCl pH8) and finally refixed for 20 minutes with a 4% PFA. Following five additional PBST washing steps, embryos were incubated at 68 °C in L1 buffer (50% De-ionized formamide; SSC 5X pH4.5; 1% SDS; 0.1% Tween 20) for 10 minutes. Next, embryos were incubated for 2 hours at 68 °C in hybridization buffer 1 (L1 Buffer, tRNA 100ug/ml, heparin 50ug/ml), followed by overnight incubation at 68 °C in hybridization buffer 1 containing 150-200ng/ml of digoxygenin probe, previously denaturalize 10 minutes at 80 °C in hybridization buffer 1. On the second day, unbound probe removal involved three washes of 30-minute at 68 °C with L1 buffer, three of L2 buffer (50% De-ionized formamide; SSC 2X pH4.5; 0.1% Tween 20), and one of 15 minutes with L3 buffer (SSC 2X pH4.5; 0.1% Tween 20), followed by 40 minutes incubation at RT. After, embryos were washed three times in TBST (TBS with 1% Tween) and pre-incubated with blocking solution (10% serum/TBST) for 2 hours, before overnight incubation at 4 °C in blocking solution containing a 1/5000 dilution of anti-digoxigenin-alkaline phosphatase. On the third day, unbound antibody was removed through a series of 30-minute washes at room temperature with TBST, followed by overnight incubation at 4 °C. On the fourth day, staining was initiated by washing at RT with NTMT solution (100mM NaCl; 100mM Tris pH9.5; 1% Tween; 50mM MgCl_2_), followed by staining with BM Purple (sigma # 11442074001). *Shox2* expression was assessed by WISH at E12.5 in *Shox2^dmCherry^*^/+^ mouse embryos. *Hoxd13* expression was assessed by WISH at E13.5 in *Shox2^Uln^*and CD1 control mouse embryos. Images were taken using in a petri dish with a top thin layer of 1% agarose on a Zeiss Axio Zoom V16 using the Axiocam 506 color camera the Zen Blue Software v3.6.

### Tissue collection and single-cell dissociation

Forelimb or Hindlimb buds of E10.5, E11.5, E12.5, E13.5 or E14.5 control (*Shox2^dmCherry/+;RosaEYFP/+^)* or mutant embryos were micro-dissected in 1X DPBS (Gibco, 14190-094) and placed in 1.5ml tubes. After DPBS removal, each tube containing pairs of limb buds were incubated with 400μl trypsin-EDTA 0.25% (Thermo Fischer Scientific, 25300062) supplemented with 40μl of 5% BSA in PBS (Sigma Aldrich, A7906-100G), during 8-9 min for small embryos (E10.5 and E11.5) or 12-15 min for larger embryos (E12.5, E13.5 or E14.5) at 37°C in a Thermomixer with a resuspension step after the first 6 min and at the end of the rest of the incubation time. After Trypsin inactivation with one volume of 5% BSA, cells were passed through a 40μm cell strainer and another volume of 5% BSA was added to wash the cell strainer. Cells were spun at 400g for 5min at 4 °C and resuspended in 1%BSA in PBS (5mM Na-Butyrate was added in case the cells were processed to be sorted and later used for downstream ChIP experiments). The single-cell suspension obtained from this process were later used for subsequent flow cytometry experiments or single-cell library preparation in the latter case cells were then counted using an automatized cell counter and a 1% BSA 700cells/ul suspension was prepared.

### Single-cell RNA-seq library preparation

Chromium Single Cell 3ʹ GEM, Library & Gel Bead Kit v3 (10X Genomics, #PN-1000075) or v3.1 (10X Genomics, #PN-1000121) was used to prepared single-cell libraries following 10X Genomics manufacturer’s protocol by the iGE3 Genomic Platform. In brief, Single-Cell 3’ Gel Beads were combined with the Master Mix containing single cells, and Partitioning Oil onto Chromium Chip B, to generate Gel Beads in Emulsion (GEMs). Full-length cDNA was produced, during beads incubation, from the poly-adenylated mRNA barcoded. Right after, gel beads were dissolved, cDNA was amplified via PCR. This cDNA was used to construct the library and then sequenced on an Illumina HiSeq 4000 or a Illumina NovaSeq 6000. On average, 7000 cells were loaded on the Chromium Chip and between 45000-58000 mean reads were obtained.

### Flow cytometry sorting and analysis

Fluorescent-activated cell sorting (FACS) was used to isolate cell populations based on the dmCherry and EYFP fluorescent signal by using the BD FACS Aria fusion with a blue laser (488nm, filter 530/30) for the EYFP signal and with a YG laser (561nm, filter 610/20) for the dmCherry signal. When sorted of cells was not required and only recording of cell population proportion was needed, we used the Beckman Coulter Cytoflex analyzer with a blue laser (488nm, filter 525/40) for the EYFP signal and with a YG laser (561nm, filter 620/20) for the dmCherry signal. In both cases, a first FSC/SCC gating was set between 30/40 and 210/240 to exclude debris followed by dead cells removal using a viability dye (DAPI, AppliChem, #A10010010 for the BDAria or DR, Invitrogen, D15106 for the Cytoflex). Then, doublets were excluded before stablishing the final gating. Flow cytometry analysis to extract to apply a homogeneous gating to the different experiments and cell proportion analysis was performed with the FlowJo™ Software (version 10.9.0). Statistical t-test for pair-wise comparison of cell proportion changes was performed using R and ggplot2 (version 3.4.4).

### RNA-seq processing and library preparation

After FACS sorting, cells were spun down at 1000g for 5 minutes at 4°C, the supernatant was removed, and then cells were snap frozen at −80°C. For each experimental condition including the frozen pellets of single-cell suspension for bulk experiment, total RNA was extracted from biological replicates containing between 3.5 × 10^4^ - 1.5 × 10^5^ cells each, using the RNeasy Micro Kit (Qiagen, 74004). Quantification of total RNA was performed with Qubit 2.0 (LifeTechnologies) and the RNA High Sensitivity Assay (Q32852). Then, libraries were prepared by the iGE3 Genomic Platform using the SMART-Seq v4 kit (Clontech, 634893) for the reverse transcription and cDNA amplification, starting from 5 ng of total RNA. For library preparation, 200 pg of cDNA were used with the Nextera XT kit (Illumina, FC-131-1096). Library molarity and quality was assessed with the Qubit and TapeStation using a DNA High sensitivity chip (Agilent Technologies). Libraries were pooled at 2 nM and loaded for clustering on a Single-read Illumina Flow cell for an average of 35 million reads/library. Reads of 50 bases were generated using the TruSeq SBS chemistry on an Illumina HiSeq 4000 or Illumina NovaSeq 6000 sequencer.

### ChIP-seq and C-HiC cell processing

After FACS sorting, cells were centrifuged at 1500g for 5 minutes at 4°C, and the supernatant was discarded. Cells were resuspended in 10% FCS/PBS and fixed either with 1% PFA (Sigma-Aldrich #252549) for ChIP-seq or with 2% PFA for C-HiC for 10 minutes rolling. Fixation was halted by adding 1.425M glycine followed by centrifugation at 1000g for 8 minutes at 4°C. Then, cells were lysed in cold lysis buffer (10mM Tris-HCl pH7.5, 10mM NaCl, 5mM MgCl_2_, 1mM EGTA, with Roche Protease Inhibitor #04693159001) and incubated on ice for 10 minutes to extract nuclei. Nuclei were then centrifugated at 1000g for 5 minutes at 4°C, washed in cold 1× PBS and centrifuged again 1000g for 1 minutes at 4°C. PBS was removed and nuclei frozen at −80°C.

### ChIP-seq immunoprecipitation and library preparation

Prior to sonication of frozen nuclei, 30ul of magnetic Protein G beads (Invitrogen 10003D) were pre-washed in 0.25%BSA/DPBS and resuspended in 1ml of L3 sonication buffer (10mM TrisCl pH 8.0, 100mM NaCl, 1mM EDTA, 0.5mM EGTA, 0.1% Na-Deoxycholate, 0.5% N-Laroylsarcosine, filtered and with Roche Protease Inhibitor #04693159001) with 1% Triton. H3K27Ac antibody was added (Diagenode, C15410174), using a 2.5ul, and beads were left to rotate at 4°C for a minimum of 4 hours.

In parallel, fixed nuclei (an average of 5 × 10^5^ cells) were sonicated 8 pulses (30 seconds ON / 30 seconds OFF) to 200–500 bp fragments using a Bioruptor Pico sonicator (Diagenode) in 200ul of L3 buffer. After sonication 1% Triton was added to the samples that were centrifuged 10 minutes at 4°C. Subsequently, chromatin was combined with the magnetic beads from which unbound antibodies were previously removed and 1.1ml of fresh L3 buffer was added. Samples were subjected to overnight rotation at 4°C. The following day unbound chromatin was removed through seven washes in RIPA buffer (1% NP-40, 0.7% Na-Deoxycholate, 1mM EDTA, 50mM HEPES-KOH, pH 7.55 and 0.5M LiCl with Roche Protease Inhibitor #04693159001) and one in TE buffer. Chromatin was then eluted and subjected to de-crosslinking overnight with the addition of 5μL Proteinase K (10mg/mL) at 65°C. This was followed by treatment with RNase A (4μL, 10mg/mL) at 37°C for 30 minutes, phenol:chloroform:IAA extraction, and precipitation (1/10 NaAc, 3ul of glycogen and 2.5 volumes of EtOH 100%) at −80°C during 30 minutes. Chromatin was then washed with 1ml EtOH 80%. Finally, the chromatin was eluted in 100μL H_2_O. Na-Butyrate (5 mM) was added to all buffers. Then, libraries were prepared by the iGE3 Genomic Platform. Briefly, ChIP-enriched DNA (<10 ng) was used to prepare libraries with the Illumina TruSeq ChIP kit, following the manufacturer’s guidelines. Libraries were validated on a Tapestation 2200 (Agilent) and a Qubit fluorimeter (Invitrogen – Thermofisher Scientific). Libraries were pooled at 2 nM and loaded for clustering on a Single-read Illumina Flow cell. Reads of 50 bases were generated using the TruSeq SBS chemistry on an Illumina HiSeq 4000 or an Illumina NovaSeq 6000 sequencer.

### C-HiC and library preparation

To prepare C-HiC libraries 1 × 10^6^ fixed frozen nuclei were used per sample. Nuclei were taken up in 520 µl of 1× DpnII buffer (NEB, R0543M) and incubated with 7.5 µl of 20% SDS for 1 hour shaking at 600 r.p.m. at 37 °C. Subsequently, 75 µl of 20% Triton X-100 was added and incubated by shaking at 600 r.p.m. at 37 °C for another hour. A 40-µl aliquot was preserved as a control for undigested chromatin (stored at −20 °C). Chromatin digestion was initiated using 600 µl of DpnII buffer and 400 Units of DpnII, shaking at 600 r.p.m. at 37 °C for 6 hours; then, 400 Units of DpnII were added, and the samples were incubated overnight, shaking at 600 r.p.m. at 37 °C. The following morning, 200 Units more of DpnII were added, and the samples were incubated 4 hours shaking at 600 r.p.m. at 37 °C. A 80-µl aliquot was extracted to assess digestion efficiency (stored at −20 °C). DpnII restriction enzyme was subsequently inactivated at 65 °C for 25 minutes. Next, the digested chromatin was diluted and religated in 5.1 ml H2O, 700 µl of 10× ligation buffer (1M Tris-HCl pH 7.5; 500mM DTT; 500mM MgCl2; 100mM ATP), and 100 Units (Weiss units) T4 DNA ligase (Thermo Fisher Scientific, #EL0013), incubated at 16 °C for 4 hours. The ligated samples were further incubated for 30 minutes at room temperature. De-crosslinking of samples and test aliquots occurred overnight by adding 30 µl and 5 µl proteinase K (10mg/ml), respectively, and incubating at 65 °C. On the following day, 30 µl or 5 µl of 10 mg/ml RNase was added to the samples and test aliquots, respectively, and incubated for 45 minutes at 37 °C. Chromatin was then precipitated by adding 1 volume of phenol-chloroform to the samples and test aliquots, vigorously shaking them, followed by centrifugation at 2200g at room temperature for 15 minutes. The upper phase containing the chromatin was transferred to a new tube. Samples were then prepared for precipitation by adding 7ml of H_2_O; 1 ml of 3M NaAc pH 5.2 and 35ml of 100% EtOH and incubated over weekend at −20 °C. The precipitated chromatin was isolated by centrifugation at 2200g for 45 minutes at 4 °C. The chromatin pellet was washed with 70% ethanol and further centrifuged at 2200g for 15 minutes at 4 °C. Finally, the 3C library chromatin pellet was dried at room temperature and resuspended in 150ul of 10 mM Tris-HCl pH 7.5. Quantification of total re-ligated product was performed using the Qubit High Sensitivity DNA Assay (Q32851). To assess the 3C library 5ul of the re-ligated sample was loaded on a 1.5% agarose gel along with the undigested and digested aliquots. Then, libraries were prepared by the iGE3 Genomic Platform. Briefly, the 3C library was then sheared using a Covaris sonicator (duty cycle: 10%; intensity: 5; cycles per burst: 200; time: 6 cycles of 60 s each; set mode: frequency sweeping; temperature: 4–7 °C). Adaptors were added to the sheared DNA and amplified according to the manufacturer’s instructions for Illumina sequencing (Agilent). Subsequently, the library was hybridized to custom-designed SureSelect beads and indexed for sequencing (50–100 bp paired-end) following the manufacturer’s instructions (Agilent). Libraries were sequenced an Illumina HiSeq 4000 or Illumina NovaSeq 6000 sequencer. *Shox2* C-HiC SureSelect library was designed using the GOPHER Java desktop application, version 0.5.7 (Hansen et al., 2019), for the genomic interval mm39:chr3:65103500-68603411 covering the *Shox2* locus and adjacent TADs.

### Custom genome for NGS analyses

All NGS datasets generated in this study were aligned to a customized version of the GRCm39/mm39 assembly (Cunningham et al., 2022) incorporating the dmCherry-P2A-CRE and floxed-SV40pASTOP-EYFP cassettes as artificial chromosomes, that we termed as GRCmm39/ mm39_dsmCherry_P2A_CRE_EYFP genome. NGS datasets downloaded from GEO and re-analyzed in this study (Andrey et al., 2017; Sheth et al., 2016) were aligned to the normal GRCm39/mm39 assembly (Cunningham et al., 2022). For annotation, GTF files sourced from ENSEMBL GRCm39 release 104 (Cunningham et al., 2022) were used, with a filtering process applied to exclude read-through/overlapping transcripts. Only transcripts annotated as ‘protein-coding’ for their respective genes were retained, while those flagged as ‘retained_intron’, ‘nonsense-mediated decay’, etc., were discarded. This filtration aimed to retain only unambiguous exons, mitigating potential quantitative biases during data analysis conducted using STAR/Cufflinks (Amandio et al., 2016).

### Single-cell analysis

#### Processing of sequenced reads

Demultiplexing, alignment, filtering of barcodes, and UMI counting of two replicates for each stage of interest E10.5, E11.5 and E12.5, except from E13.5 that only one replicate was produced, were executed using the 10× Genomics Cell Ranger software (version 6.1.2) in accordance with the manufacturer’s guidelines, default settings and custom genome GRCmm39/ mm39_dsmCherry_P2A_CRE_EYFP built following using the cellranger mkref pipeline. Cell Ranger output files for each dataset were further processed using the velocyto run10x command from the velocyto.py tool (version 0.17.17) (La Manno et al., 2018) in Python (version 3.9.12) with our custom genome and the UCSC genome browser repeat masker.gtf file to mask expressed repetitive elements to generate a loom file for each sample. Each resulting loom matrix, comprising spliced/unspliced/ambiguous reads, was individually imported into R (version 4.1.2) using the Read Velocity function from the Seurat Wrappers package (version 0.3.0). Simultaneously, feature-filtered output matrices obtained from Cell Ranger were loaded into R separately through the Read10X function of the Seurat package (version 4.2.1) (Stuart et al., 2019). Subsequently, the spliced, unspliced, ambiguous, and RNA feature data were combined into a single matrix for each dataset. Following this, each matrix was transformed into a Seurat object using the Seurat package. Consequently, for each sample, a single Seurat object was obtained, encompassing four assays. Three of these assays (spliced, unspliced, and ambiguous) were used for downstream RNA velocities estimations, while the RNA feature assay was employed for subsequent gene expression analysis among the samples, as detailed below.

#### Quality control and filtering

Quality control and pre-processing of each Seurat object for our samples were conducted based on the following criteria. Cells expressing fewer than 200 genes or exhibiting more that 7500 features were excluded from the analysis. Additionally, we calculated the proportion of reads mapping to the mitochondrial genome, filtering out cells with a mitochondrial content exceeding 5%, as elevated levels of mitochondrial mRNA have been linked to cell death. Conversely, cells with a mitochondrial content lower than 0.5% were also excluded, as our observations suggest that these cells likely originate from blood cells, possibly due to the dissection protocol. After this step we decided to continue the analysis with only one replicate per stage. The replicate with the best quality control for each stage was selected.

#### Individual dataset normalization, scaling, and dimensional reduction

After filtering each dataset was individually normalized using the default parameters provided by Seurat for the LogNormalize method and applying it to the RNA features assay. Subsequently, we calculated the most variable features excluding the *CRE*, *EYFP* and *dmCherry* artificial genes added on our custom genome from the list of variable genes to avoid that they drive the PCA. Then, scaling was performed via linear transformation and scaled data were then employed for principal component analysis (PCA), utilizing the default 50 principal components (PCs). Additionally, non-linear dimensional reduction was conducted using Uniform Manifold Approximation Projection (UMAP) (Leland McInnes et al., 2018), with 1:50 dimensions utilized as input.

#### Cell doublet identification and features annotation

Pre-processed and normalized datasets were individually examined to detect putative doublet cells. Doublets identified in each dataset were subsequently excluded using the DoubletFinder R package (version 2.0.3) (McGinnis et al., 2019). The doublet rate (nExp parameter) utilized was estimated based on the number of cells captured and pK parameter was estimated following the strategy defined in the package, resulting in the following values: *Shox2^trac^* Hindlimb E10.5, nExp= 89, pK=0.3; E11.5 nExp= 98, pK=0.16; E12.5 nExp= 71, pK=0.25; E13.5 nExp= 88, pK=0.1. After removing doublets, counts for *CRE*, *dmCherry*, *Shox2*, and *EYFP* per cell were estimated. Given the low counts for *dmCherry* (likely a limitation due to the 10X single-cell technique where transcripts are only sequenced from the 3’ poly-A end, which does not allow for adequately cover the *dmCherry* sequence), we proceeded with further cell classification using only *Shox2*, *EYFP*, and *CRE* counts. Moreover, from that point on, we referred to *CRE* counts as *dmCherry-P2A-CRE*, since we assumed that could be used as a proxy for both *dmCherry* and *CRE* genes. Of note, due to the shared SV40polyA tail between *dmCherry-P2A-CRE and EYFP*, many reads were ambiguous, leading to a limited number of reads assignable to *dmCherry-P2A-CRE*, which we anticipated to be lower in expression than *EYFP* (as constitutively expressed from the *ROSA26* promoter). Cells were then classified as positive for each of these genes if they had at least one count, and negative otherwise. Cells positive for *Shox2*, negative for *EYFP*, and either positive or negative for *dmCherry-P2A-CRE* were classified as *initiating*. Those positive for both *Shox2* and *EYFP*, regardless of *dmCherry-P2A-CRE* status, were classified as *maintaining*. Cells negative for both *Shox2* and *dmCherry-P2A-CRE* but positive for *EYFP* were considered *decommissioned*. Cells negative for all three genes were marked as inactive, and any remaining combinations of gene expression fell under the class “other”. A new metadata column containing the classification of the cells was then created.

#### Merge of all datasets and normalization

All datasets were then merged into a single Seurat object without undergoing integration, allowing for ensemble downstream analysis of the four datasets. Subsequently, no batch effect was observed in this merged dataset. A new column of the metadata was created at this step to label samples based on the stage, to keep this information for downstream analysis. Afterward, we applied the SCTransform normalization protocol (Hafemeister and Satija, 2019) to our newly merged Seurat object, utilizing default parameters, over the spliced assay.

#### Cell-cycle scoring and cell-cycle and stage regression

Since we observed, during individual dataset analysis, that a portion of the variance was attributable to cell-cycle genes, we assigned cell cycle score using the CellCycleScoring function implemented in Seurat. As we also observed that sample variance by a stage effect we regress out the cell-cycle heterogeneity and stage variability by applying SCTransform normalization method to our merged object, using the spliced assay as the source, and incorporating the calculated cell-cycle scores (S.Score and G2M.Scores) and the stage metadata information as variables to regress, in addition to the default settings. Subseqeuntly, we excluded *dmCherry*, *CRE* and *EYFP*, if they were present from the variable genes to avoid that they drive the PCA.

#### Clustering

Following the regression step, cells were clustered using the standard steps of the SCTransform Seurat workflow. Briefly, PCA (npcs = 50), UMAP (dims = 1:50, n.neighbors = 50), and nearest neighbors were calculated. Clusters were identified using the Seurat FindClusters function with default parameters and a resolution of 0.7, resulting in the definition of 21 clusters. Cluster identity was determined by assessing the expression difference of each gene between each cluster and the rest of the clusters using the FindMarkers function. Clusters presenting similar features profiles were combined, reducing the final number of clusters identified to 6 clusters (**Supplementary Fig. S2A)**. The mesenchyme (comprising 13 out of the 21 clusters), epithelium (consisting of 3 out of 21), muscle (comprising 2 out 21), and endothelium, immune Cells, and blood Cells clusters each represented by only 1 cluster. The presence of expected identity markers in the new clustering was confirmed by running the FindMarkers function with default parameters and using grouping.var = “stage” and only.pos = TRUE.

#### Subsetting and re-clustering

Given the focus of this study on populations expressing *Shox2*, we subsetted and re-clustering the mesenchyme cluster. To do, after applying subset function to the “Mesenchyme” UMAP embedding was computed with the following parameters: dims = c(1:10), n.neighbors = 30L, min.dist = 0.5, metric = “euclidean”, spread = 1, while keeping all other parameters at their default values. Subsequently, the cluster resolution after finding neighbors was set at 1.1 to reveal subpopulations. We observed 18 mesenchyme subpopulations, each named based on their identity genes. Identity markers were identified using the FindMarkers function on the RNA assay, with grouping.var = “stage”, only.pos =TRUE, logfc.threshold = 0.3, min.diff.pct = 0.1, and all other parameters set to default values. Clusters presenting similar features profiles were combined, reducing the final number of clusters identified to 15 clusters (**Fig. 2B**), late proximal progenitors (comprised 3 out of the 18 clusters) and irregular connective tissue (contained 2 out the 18) the other clusters remained represented by 1 cluster. Final identity markers for the new clustering was assessed by running the FindMarkers function on the RNA assay, only.pos =TRUE, logfc.threshold = 0.5, pseudocount.use = 0, min.diff.pct = 0.1, and all other parameters set to default values (**Supplementary Table S2**).

#### RNA-velocity analysis

For the RNA-velocity analysis, we used the unspliced (immature) and spliced (mature) abundances calculated for each replicate of our datasets, as described earlier (see Material and Methods, Single-cell analysis, Processing of sequenced reads). We then performed RNA-velocity analysis on all combined datasets by exporting Seurat object as h5Seurat files using the SeuratDisk package (version 0.0.0.90) and for using it as input in Scvelo (version 0.2.5) (Bergen et al., 2020) in Python (version 3.9.16). Then the standard protocol described in scVelo was followed, with the exception of using npcs = 10 and n.neighbors = 30, to match the parameters used for UMAP embedding in Seurat. *Shox2* velocity was computed by running velocyto package for R (version 0.6) (La Manno et al., 2018) with default parameters on the Seurat Object to generate an embedding file from which *Shox2* was only plotted using ggplot2 (version 3.4.4).

#### Graphical plots

FeaturePlot and VlnPlot were generated from the RNA assay of the Seurat objects. FeaturePlot and Dimplot were produced using default Seurat parameters. Density UMAP plots were produced using the Nebulosa v1.4.0 package (Alquicira-Hernandez and Powell, 2021). Cell proportions were calculated using the prop.table tool from the base R package (version 4.1.2) followed by plotting using ggplot2 (version 3.4.4).

### RNA-seq analysis

#### RNA-seq reads processing

FASTQ files from FACS-sorted cells or entire limbs generated in this study were processed using CutAdapt v1.18 to trim NextSeq adapter sequences and low-quality bases (Martin, 2011), employing the adapter sequence -a CTGTCTCTTATACACATCTCCGAGCCCACGAGAC with a quality cutoff of 30 (-q30) and a minimum length requirement of 15 bases (-m15). In the case of the samples from GEO datasets that we wanted to reanalyze (Andrey et al., 2017) CutAdapt was used to trim TruSeq adapter sequences and low-quality bases, using the following parameters -a GATCGGAAGAGCACACGTCTGAACTCCAGTCAC, -q30 and -m15). Unstranded reads were then mapped on the customized genome GRCm39/ mm39_dsmCherry_P2A_CRE_EYFP, in the case of the datasets produced in this study, or to the GRCm39/mm39, in the case of the reanalyzed dataset. The STAR version 2.7.2b (Dobin et al., 2013) was then used together with the filtered GTF file generated for this study (see Custom genome for NGS analyses in this Material and Methods section) for accurate gene quantification using tailored settings (--outSAMstrandField intronMotif--sjdbOverhang ‘99’ -- sjdbGTFfile $gtfFile--quantMode GeneCounts--outFilterType BySJout--outFilterMultimapNmax 20 -- outFilterMismatchNmax 999--outFilterMismatchNoverReadLmax 0.04--alignIntronMin 20 -- alignIntronMax 1000000--alignMatesGapMax 1000000--alignSJoverhangMin 8 -- alignSJDBoverhangMin 1). FPKM values were then determined by Cufflinks version 2.2.1 (Roberts et al., 2011) using the filtered GTF file generated for this study (see Custom genome for NGS analyses in this Material and Methods section) and tailored settings (--max-bundle-length 10000000 -- max-bundle-frags 100000000 -- multi-read-correct--library-type “fr-firststrand”--no-effective-length-correction -M MTmouse.gtf). Then, Normalized FPKM were computed by determining coefficients extrapolated from a set of 1000 house-keeping genes stably expressed across the series of compared RNA-seq datasets (Brawand et al., 2011). Differential expression analysis was performed using DEseq2 (Love et al., 2014) R package (version 1.34.0) with the Wald test for comparisons across samples and multiple test correction using the FDR/Benjamini-Hochberg test.

### ChIP-seq analysis

#### ChIP-seq reads processing

Reads from ChIP-seq sequencing either from dataset generated for this study or from GEO datasets that we wanted to reanalyze (Andrey et al., 2017; Sheth et al., 2016) were processed first using CutAdapt version 1.18 (Martin, 2011) to trim TruSeq adapter sequences and low-quality bases, specifying the adapter sequence with -a GATCGGAAGAGCACACGTCTGAACTCCAGTCAC, a quality threshold of 30 with -q30, and a minimum length of 15 bases with -m15. Then reads were mapped to our GRCm39/mm39_dsmCherry_P2A_CRE_EYFP customized mouse genome, in the case of the datasets generated for this study, or to the GRCm39/mm39, in the case of the reanalyzed datasets using Bowtie2 version 2.3.5.1 (Langmead and Salzberg, 2012) with its default settings.

Subsequently, only reads with a mapping quality score (MAPQ) of 30 or higher were retained, as filtered with SAMtools view version 1.10 (Danecek et al., 2021). For coverage and peak analysis, reads were extended by 200 base pairs and processed using MACS2 version 2.2.7.1 (Zhang et al., 2008) with the parameters --broad --nolambda --broad-cutoff 0.05 --nomodel --gsize mm --extsize 200 -B 2 for broad peak calling, in the case of H3K27ac ChIP produced from the FACS sorted cells for this study. For reanalyzed datasets coverage and peak analysis, reads were extended by 200 base pairs and processed using MACS2 version 2.2.7.1 (Zhang et al., 2008) with the parameters --call-summits -- nomodel --extsize 200 -B 2 for narrow peak calling. The coverage normalization was performed by MACS2, adjusting for the total millions of tags used in the analysis.

#### ChIP-seq reads visualization

For the reprocessed *Hoxa13* and *Hoxd13* ChiP-seq (Sheth et al., 2016), when a peak with a score >100 (MACS2) overlap one of our enhancers, we considered the region bound.

#### Early, common, late putative enhancers classification on entire forelimb datasets

RNA-seq FASTQ files from two replicates of entire forelimbs at E10.5 and E13.5 (Andrey et al., 2017) were re-analyzed following the RNA-seq pipeline previously described (see RNA-seq analysis in this Material and Methods section). Genes related to limb development were selected for downstream analysis. Average of normalized FPKM values was calculated and used to compute the ratio among E10.5 and E13.5 datasets. Then, since we were interested in genes having stable expression between E10.5 and E13.5, they were filtered to keep those with FPKM values bigger than 5, at both stages, and we excluded genes having a fold change larger than 3 between the two stages. By applying this filtering 90 genes were selected (**Supplementary Table S1**). To analyze putative enhancers ChIP-seq H3K27Ac datasets of entire forelimbs at E10.5 and E13.5 (Andrey et al., 2017) were first reanalyzed following the ChIP-seq pipeline previously described in this material and methods section. H3K27ac MACS2 narrowpeaks were then restricted within the interaction domain defined by promoter Capture-C (Andrey et al., 2017) of the 90 filtered genes, using bedtools (version v2.30.0) intersect function (Quinlan and Hall, 2010). Then, H3K27Ac peaks around gene promoters were excluded by filtering against a −2kb/500bp window centered at the transcription start site of coding genes using again bedtools intersect. Remaining peaks were extended by +/- 300bp, using bedtools slope and merge function (Quinlan and Hall, 2010). H3K27Ac peaks were then classified as putative common enhancers when present in both E10.5 and E13.5 using bedtools intersect. H3K27Ac peaks present only in the E10.5 dataset were classified as putative early enhancers while H3K27Ac peaks present only in the E13.5 dataset were classified as putative late enhancers. Putative enhancers were then assigned to gene interaction domain (**Supplementary Table S1**). In those cases, where putative enhancers were within the overlapping region of two domains putative enhancers were assigned to the two loci.

#### Early, common, late putative enhancers classification on FACS sorted maintaining forelimb datasets

MACS2 BroadPeak files from FACS sorted maintaining cells from forelimbs at E10.5, E11.5, E12.5 and E13.5 were used to build bed files. These files were used to merge peaks within 600bp of each other using bedtools (Quinlan and Hall, 2010). Then, bedops (version 2.4.41) (Neph et al., 2012)--merge operation was used to flatten all disjoint, overlapping, and adjoining element regions into contiguous, disjoint regions peaks among the four different stages. Subsequently, peaks were extended by +/- 300bp, using bedtools slope function. Since, we wanted only to explore the putative enhancers of *Shox2* locus using bedtools intersect we selected only the region with the following coordinates mm39 chr3:65,885,132-67,539,263, in that way we created a list of peaks of interest. Peaks falling on gene promoters were manually excluded. Then as we wanted to stablish our new early, late, common enhancer classification having into consideration the scores assigned to each peak for each stage analyzed, we used deeptools (Ramirez et al., 2016) multiBigwigSummary function to compute the average scores for each peak in our curated list at each stage. Subsequently, peaks with a coverage lower than 0.3 and peaks smaller that 600bp were excluded. Finally, we analyze the slope of H3K27ac coverage across the four stages and we classified enhancers as early (<-0.6), common (>-0.6, <0.6), or late (>0.6).

### C-HiC analysis

The preprocessing and alignment of paired-end sequencing data, along with the filtering of mapped di-tags, were conducted using HiCUP pipeline (version 0.6.1) (Wingett et al., 2015) using default parameters for the configuration file and adding Nofill: 1 parameter. Bowtie2 (version 2.3.4.2)(Langmead and Salzberg, 2012) was used by the pipeline for mapping. Subsequently, filtered di-tags were processed with Juicer Tools (v1.9.9) (Durand et al., 2016) to generate binned contact maps (5kb and 10kb) from valid and unique reads pairs with MAPQ≥30 and normalized maps using Knights and Ruiz matrix balancing (Knight and Ruiz, 2013). For binning and normalization, only the genomic region mm39:chr3:65103500-68603411 covering the *Shox2* locus and adjacent TADs was considered. Subtraction maps were produced from the KR normalized maps and scaled together across their subdiagonals. CHiC maps of count values, as well as subtraction maps, were visualized as heatmaps in which values above the 99-th percentile were truncated for visualization purposes.

## Data availability

Sequencing data are available in the GEO repository under the accession number GSE262006.

## Acknowledgements

We thank Mylène Docquier, Brice Petit, Didier Chollet and Christelle Barraclough from the iGE3 sequencing facility. We thank Grégory Schneiter, Lan Tran and Cécile Gameiro from the Flow Cytrometry facility. We thank Olivier Fazio, Angélique Vincent and Fabrizio Thorel from the Transgenic facility. We thank Lucille Delisle for bioinformatic support. We thank Nicolas Liaudet from the Bioimaging facility and Stéphane Pàges, Laura Batti and Ivana Gantar from Advanced Light Sheet Imaging Center (ALICe) at the Wyss Center for Bio and Neuroengineering, Geneva. We thank Frank Costantini and Shankar Srinivas for sharing the map and plasmid sequence of their R26R-YFP targeting construct. We thank John Cobb for providing the plasmid for the *Shox2* probe. We thank the Duboule lab for discussions. The computations were performed at University of Geneva using Baobab HPC service. This study was supported by grants from the Swiss National Science Foundation PP00P3_176802, PP00P3_210996.

## Author contributions

G.A. conceived the project. R.R.G. designed the targeting constructs and deletion experiments. R.R.G. and G.A performed embryo imaging. R.R.G., A.R. and O.B. performed single dissociation and fixation for scRNA-seq, ChIP-seq and C-HIC. R.R.G. performed the scRNA-seq, RNA-seq ChIP-seq, C-HIC and associated bioinformatic analyses. F.D. performed the limb-wide analysis of early-late enhancers. R.R.G. and A.R. performed mESC targetings and prepared the cells for tetraploid aggregation. G.S. cloned and targeted the Rosa26 EYFP construct in mESCs. A.R. performed the WISH. G.A. and R.R.G. wrote the manuscript with input from the remaining authors.

## Competing interests

The authors declare no competing interests.

## Notes

### Competing Interest Statement

The authors have declared no competing interest.

