## Supplementary Figures 1-11 for "Temporal constraints on enhancer usage shape the regulation of limb gene transcription"

### Supplementary Figure S1

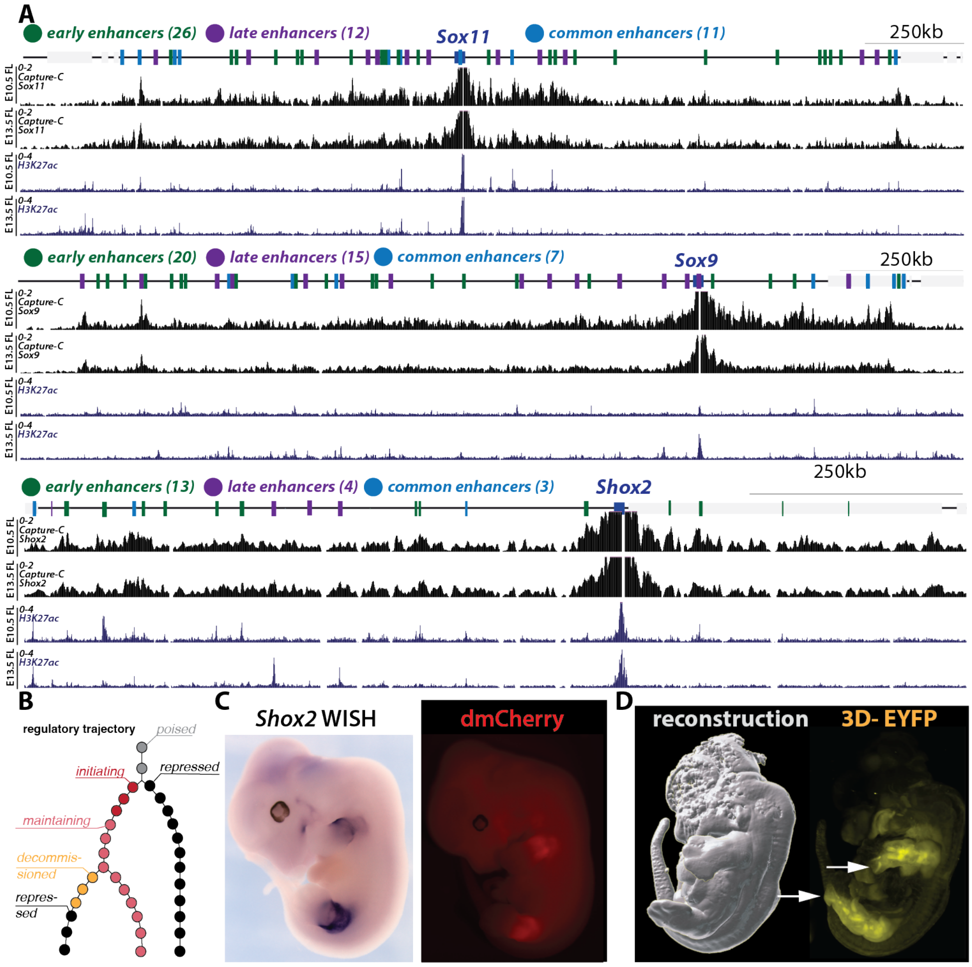


***Supplementary Figure S1. A.*** *Capture-C interaction profiles (from promoters) and H3K27Ac ChIP profile for examples of developmental loci: Sox11 (contact domain shown, mm39; chr12:26470941-28552347), Sox9 (contact domain shown, mm39; chr11:111354041-113105934), Shox2 (contact domain shown, mm39; chr3:66194862-67308968) with early (green balls), common (light blue balls), and late (purple balls) putative enhancer regions based on re-analyzed from (Andrey et al., 2017).* ***B.*** *A hypothetical regulatory trajectory begins from a poised state (grey) either towards an inactive state, therefore to locus repression (black) or towards and active state thereby to gene transcriptional initiation (dark red), followed by either transcriptional maintenance (light red) or decommissioning (orange). Eventually, repression (black) can shut down the locus.* ***C.*** *Shox2 WISH and dmCherry fluorescence in a Shox2$^{dmCherry/+}$ E12.5 embryo.* ***D****. Light sheet microscopy reconstruction of EYFP signal in a Shox2$^{trac}$ (Shox2$^{dmCherry/+}$; Rosa26$^{loxEYFP/+}$) E12.5 embryo. Note the EYFP signal in digit condensations (white arrows).*

**Supplementary Figure S2**

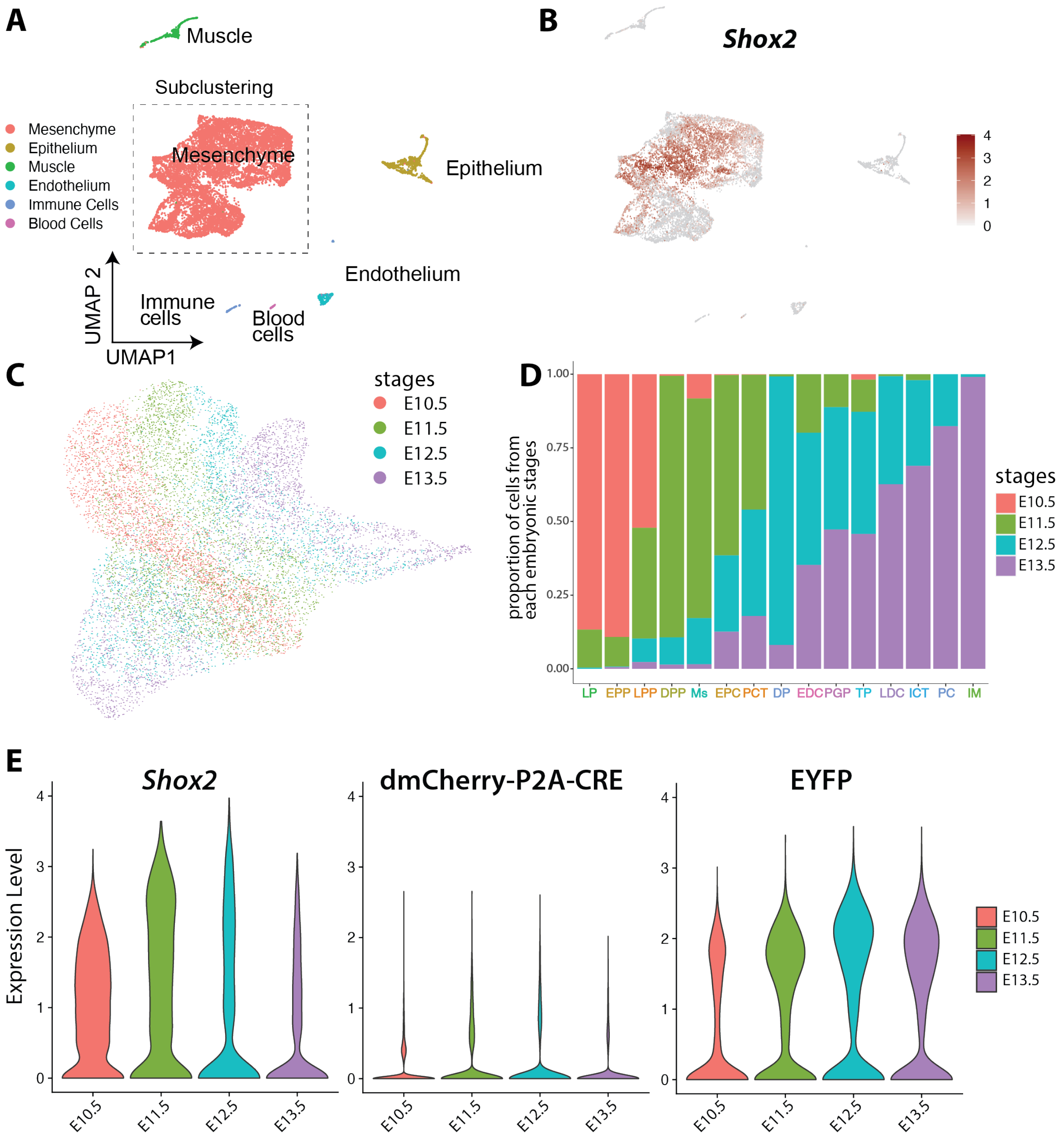


***Supplementary Figure S2: A.*** *UMAP clustering of Shox2$^{trac}$ E10.5, E11.5, E12.5 and E13.5 hindlimb cells shows one mesenchymal cluster containing most cells as well as five non-mesenchyme satellite clusters.* ***B.*** *Expression of Shox2 across all limb cell types.* ***C.*** *UMAP representation of mesenchyme split by development stages.* ***D.*** *Proportion of cell split by developmental stage in each mesenchymal cluster, ordered from early to late development.* ***E.*** *Violin plot of Shox2, dmCherry-P2A-CRE expression across stages and mesenchyme.*

**Supplementary Figure S3**

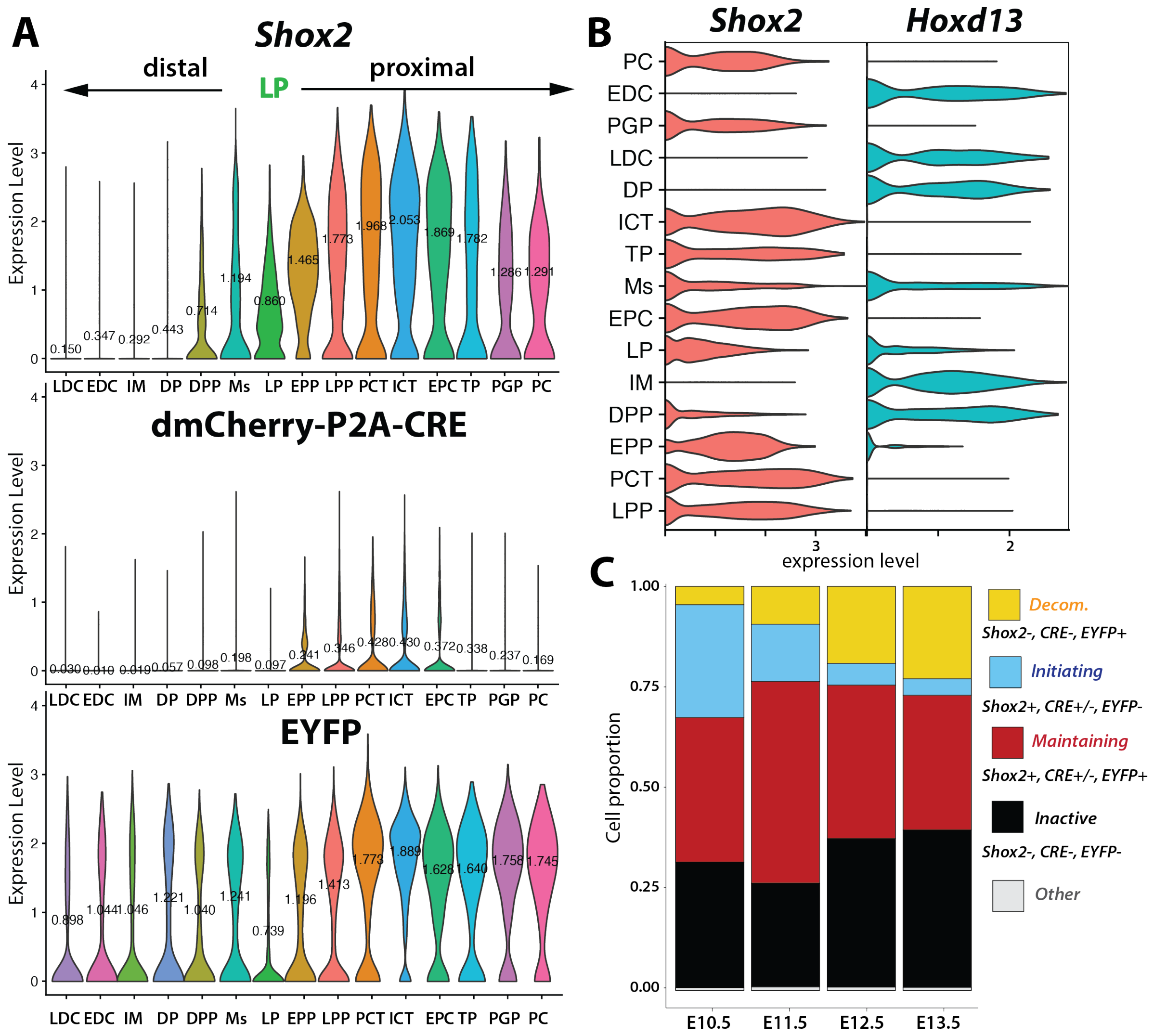


***Supplementary Figure S3: A.*** *Expression of Shox2, dmCherry-P2A-CRE and EYFP in each mesenchymal cluster, ordered according to their distal and proximal identity and to developmental time.* ***B.*** *Expression of Shox2 and Hoxd13 in each mesenchymal cluster. Note the mutual exclusion of both transcripts except for LP and Ms, and to a smaller extent for DPP.* ***C.*** *Distribution of each Shox2 transcriptional phases: initiation (Shox2+, dmCherry-P2A-CRE +/-, EYFP-, light blue), maintaining (Shox2+, dmCherry-P2A-CRE +/-, EYFP+, red), decommissioning (Shox2-, dmCherry-P2A-CRE-, EYFP+, yellow), inactive (Shox2-, dmCherry-P2A-CRE-, EYFP-, black), or other cells (when where not included in any of the previously mentioned class) across developmental stages in all cells including non-mesenchyme clusters.*

### Supplementary Figure S4

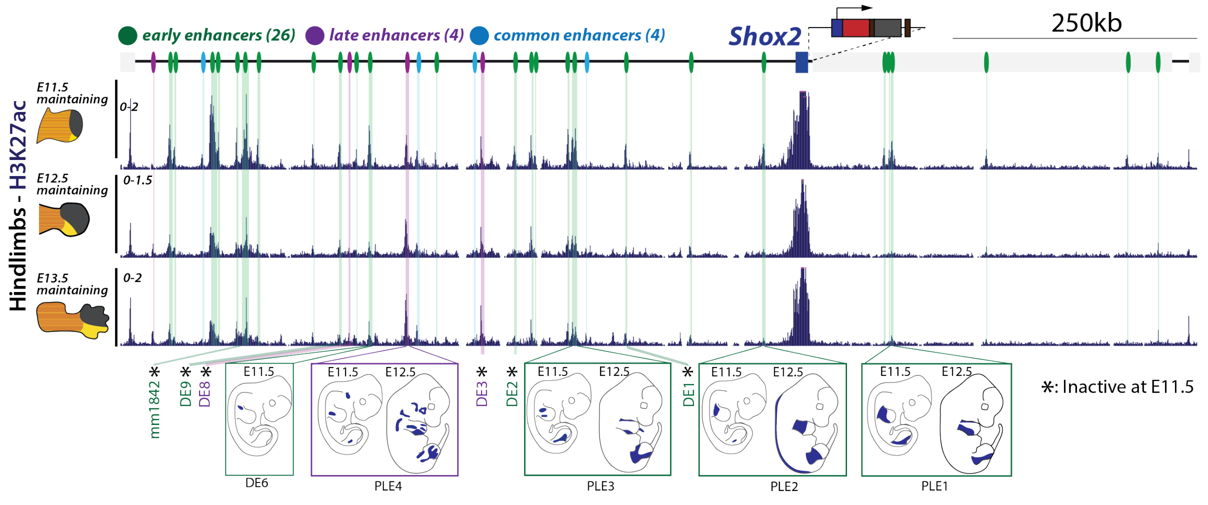


***Supplementary Figure S4:*** *H3K27ac ChIP-seq profiles of FACS sorted maintaining (dmCherry+/EYFP+) cells across E11.5, E12.5, and E13.5 hindlimbs (mm39: chr3:66,190,000-67,290,000). Putative enhancers are delineated by color-coded lines: green for early, light blue for common, and purple for late enhancers, as detailed* ***in Supplementary Table S4****. Bottom part shows a schematic representation of the pattern displayed by enhancers previously validated through in vivo LacZ reporter assays (Abassah-Oppong et al., 2023; Osterwalder et al., 2018).*

### Supplementary Figure S5

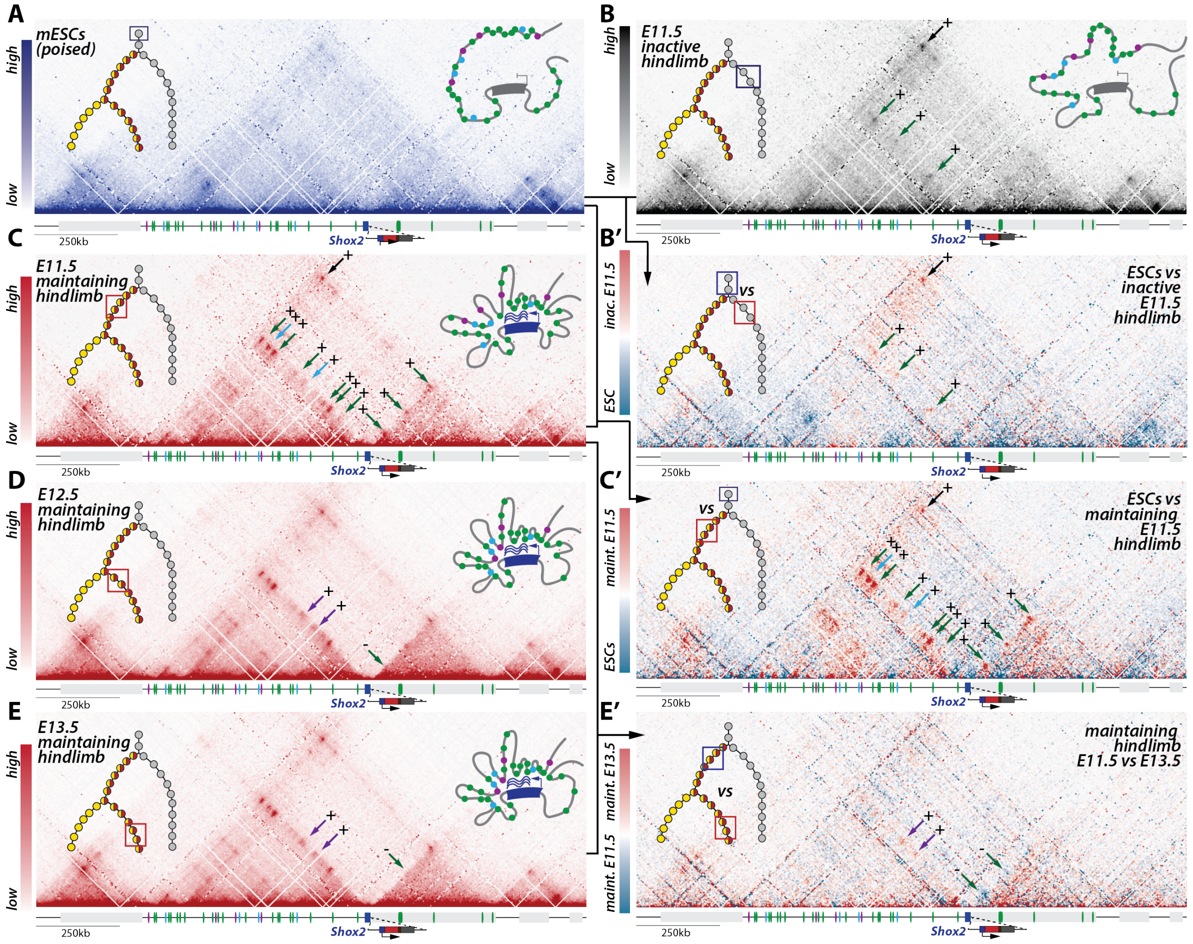


**Supplementary Figure S5:** ***Shox2 locus 3D topology associates with active enhancer-promoter interactions in hindlimbs.*** *In all Capture-HiC (C-HiC) maps (mm39: chr3:65,885,132-67,539,263), the upper left illustration represents the position of the investigated cells in the regulatory trajectory and the upper right one a model of the average 3D locus structure. The light grey box next to Shox2 is the Rsrc1 gene.* ***A.*** *C-HiC maps of the Shox2 locus in Shox2*$^{trac}$ *mESCs. Note a large TAD with few focal interaction points.* ***B.*** *C-HiC maps of the Shox2 locus in E11.5 hindlimb FACS-sorted inactive cells. Note the formation of specific contacts with three early enhancers (green arrows) and increased loop contact between the two TAD borders (black arrow).* ***B'.*** *Subtraction C-HiC map between Shox2*$^{trac}$ *E11.5 hindlimb FACS-sorted inactive cells and Shox2*$^{trac}$ *mESC.* ***C-E.*** *C-HiC maps of the Shox2* ***C.*** *E11.5* ***D.*** *E12.5, and* ***E.*** *E13.5 hindlimb FACS-sorted maintaining cells.* ***C'.*** *C-HiC subtraction maps between Shox2*$^{trac}$ *mESCs and Shox2*$^{trac}$ *E11.5 hindlimb FACS-sorted maintaining cells.* ***E'*** *C-HiC subtraction maps between E12.5 and E13.5 Shox2*$^{trac}$ *FACS sorted hindlimb maintaining cells. Changes in enhancer-Shox2 interactions are marked by colored arrows at each stage: green for early enhancers, purple for late enhancers, and light blue for common enhancers. A plus sign (+) denotes a gain of interaction, and a minus sign (-) indicates a loss of interaction relative to the previous stage. Also note the increased separation between the two subTADs at the position of the Shox2 gene body. Maps coordinates mm9; chr3:65,781,633-67,435,852. Maint. = maintaining; inac. = inactive.*

### Supplementary Figure S6

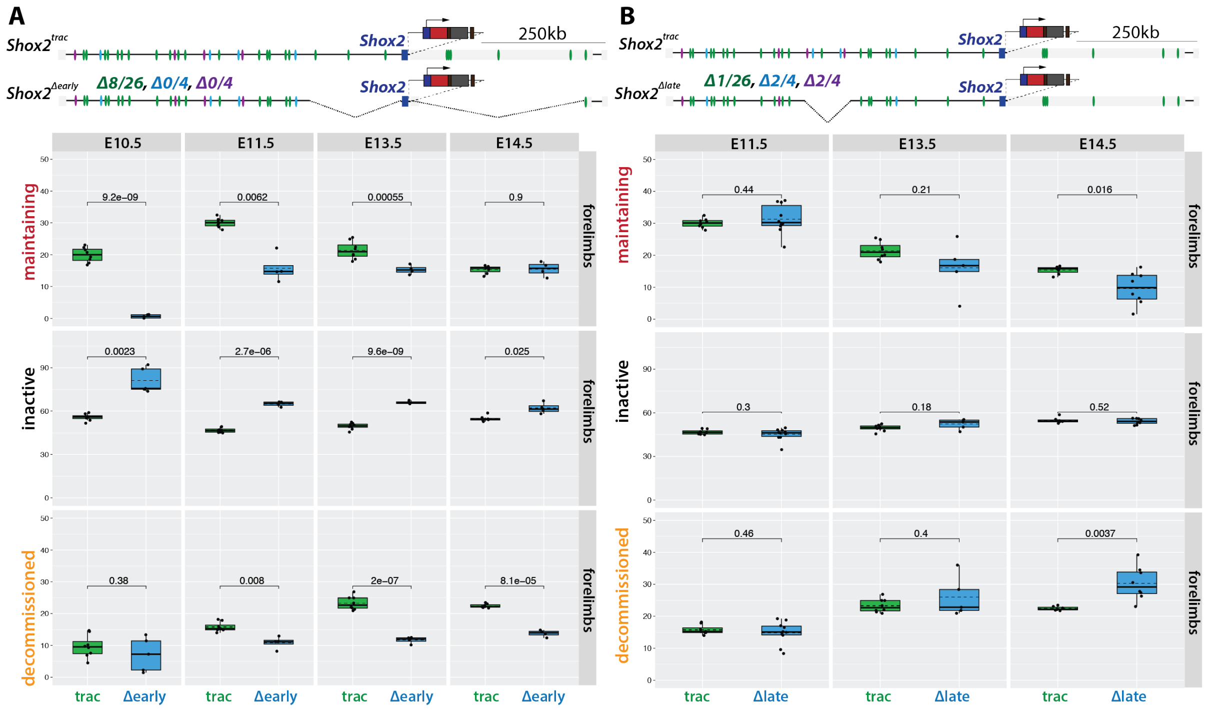


***Supplementary Figure S6. A.*** *Boxplot representation of maintaining, inactive, and decommissioned cell populations identified by flow cytometry analysis in* $Shox2^{trac}$ *versus* $Shox2^{\Delta early}$ *forelimbs at E10.5, E11.5, E13.5, and E14.5. Statistical significance was assessed using T-tests on replicates. Upper schemes illustrate the* $Shox2^{\Delta early}$ *deletion allele lacking 8 out of 26 putative early enhancers (in green), while late (in purple) and common (light blue) putative enhancers remain intact. Each dot represents one replicate.* ***B.*** *Boxplot representation of maintaining, inactive, and decommissioned cell populations identified by flow cytometry analysis in* $Shox2^{trac}$ *versus* $Shox2^{\Delta late}$ *forelimbs at E11.5, E13.5, and E14.5. Statistical significance was assessed using T-tests on replicates. Upper schemes illustrate schematic representation of the* $Shox2^{\Delta late}$ *deletion allele lacking 2 out 4 late (in purple), 2 out of 4 common (in light blue), 1 out of 26 early putative enhancers. Each dot represents one replicate.* ***A-B****, number of replicates for each forelimb genotype:* $Shox2^{trac}$ *N at* E10.5= 8, *N at* E11.5 = 8, N at E13.5 = 8, N at E14.5 = 7; $Shox2^{\Delta early}$ *N at* E10.5=5 , *N at* E11.5 = 4, N at E13.5 = 4 , N at E14.5 = 4; $Shox2^{\Delta late}$, *N at* E11.5 = 10, N at E13.5 = 5, N at E14.5 = 8.

### Supplementary Figure S7

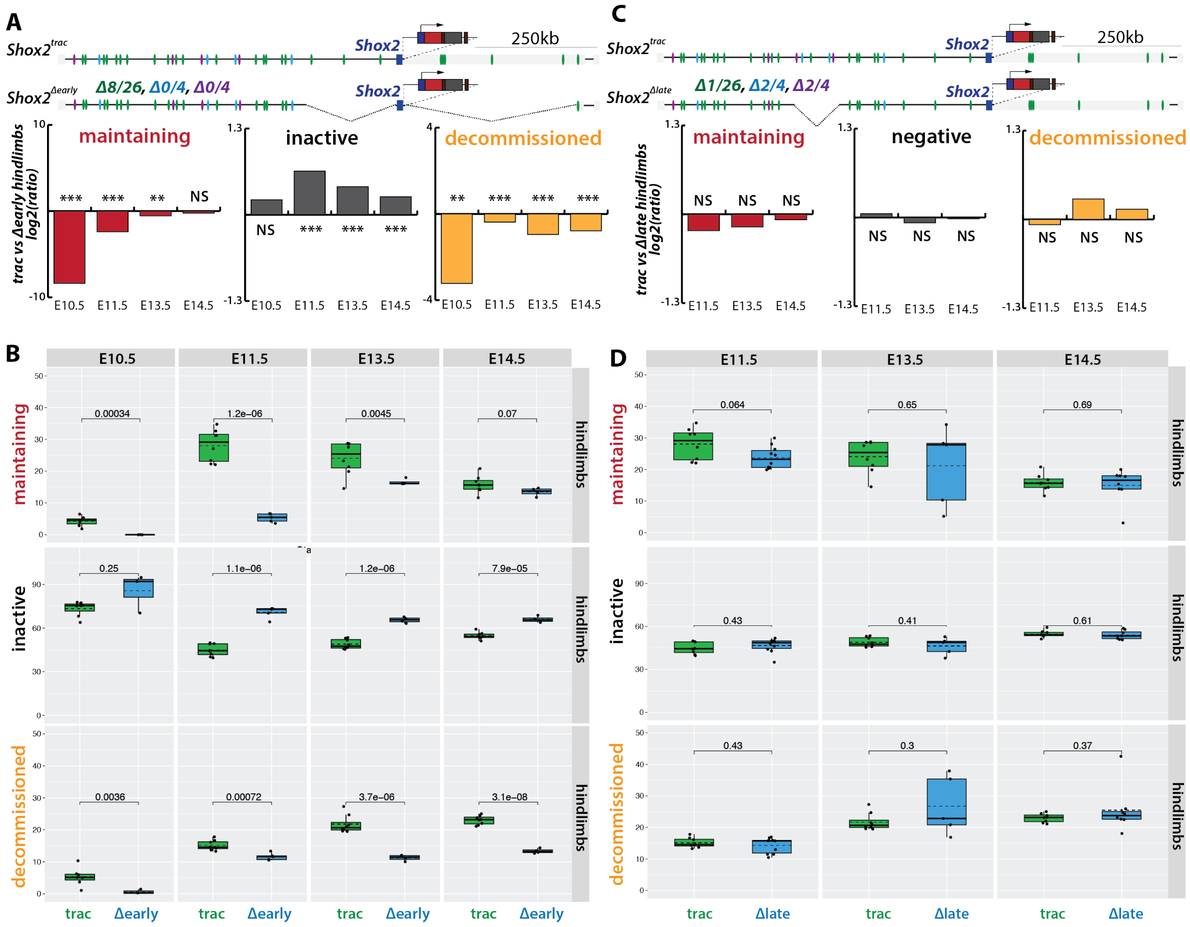

**Supplementary Figure S7:** ***A.*** *Upper part of the panel illustrates Shox2$^{\Delta early}$ deletion allele lacking 8 out of 26 putative early enhancers (in green), while late (in purple) and common (light blue) putative enhancers remain intact. Bottom part shows log2 ratio of the proportion of maintaining, inactive, and decommissioned cell populations, identified by flow cytometry analysis, in Shox2$^{trac}$ versus Shox2$^{\Delta early}$ hindlimbs at E10.5, E11.5, E13.5, and E14.5. NS= non-significant, *=p<0.05, **=p<0.01 and ***=p<0.001.* ***B.*** *Boxplot representation of maintaining, inactive, and decommissioned cell populations, identified by flow cytometry analysis, in Shox2$^{trac}$ versus Shox2$^{\Delta early}$ hindlimbs at E10.5, E11.5, E13.5, and E14.5. Statistical significance was assessed using T-tests on replicates. Each dot represents one replicate.* ***C.*** *Upper part of the panel illustrates schematic representation of the Shox2$^{\Delta late}$ deletion allele lacking 2 out 4 late (in purple), 2 out of 4 common (in light blue), 1 out of 26 early putative enhancers. Bottom part shows log2 ratio of the proportion of maintaining, inactive, and decommissioned cell population, identified by flow cytometry analysis, in Shox2$^{trac}$ versus Shox2$^{\Delta late}$ hindlimbs at E11.5, E13.5, and E14.5. NS= non-significant, *=p<0.05, **=p<0.01 and ***=p<0.001.* ***D.*** *Boxplot representation of maintaining, inactive, and decommissioned cell populations identified by flow cytometry analysis in Shox2$^{trac}$ versus Shox2$^{\Delta late}$ hindlimbs at E11.5, E13.5, and E14.5. Statistical significance was assessed using T-tests on replicates. Each dot represents one replicate.* ***B-D****, number of replicates for each hindlimb genotype: Shox2$^{trac}$ N at* E10.5= 7, *N at* E11.5 =8, N at E13.5 = 8, N at E14.5 = 8; *Shox2$^{\Delta early}$ N at* E10.5 = 3 , *N at* E11.5 = 5, N at E13.5 = 4, N at E14.5 = 4; *Shox2$^{\Delta late}$*, *N at* E11.5 = 10, N at E13.5 = 5, N at E14.5 = 8.

### Supplementary Figure S8

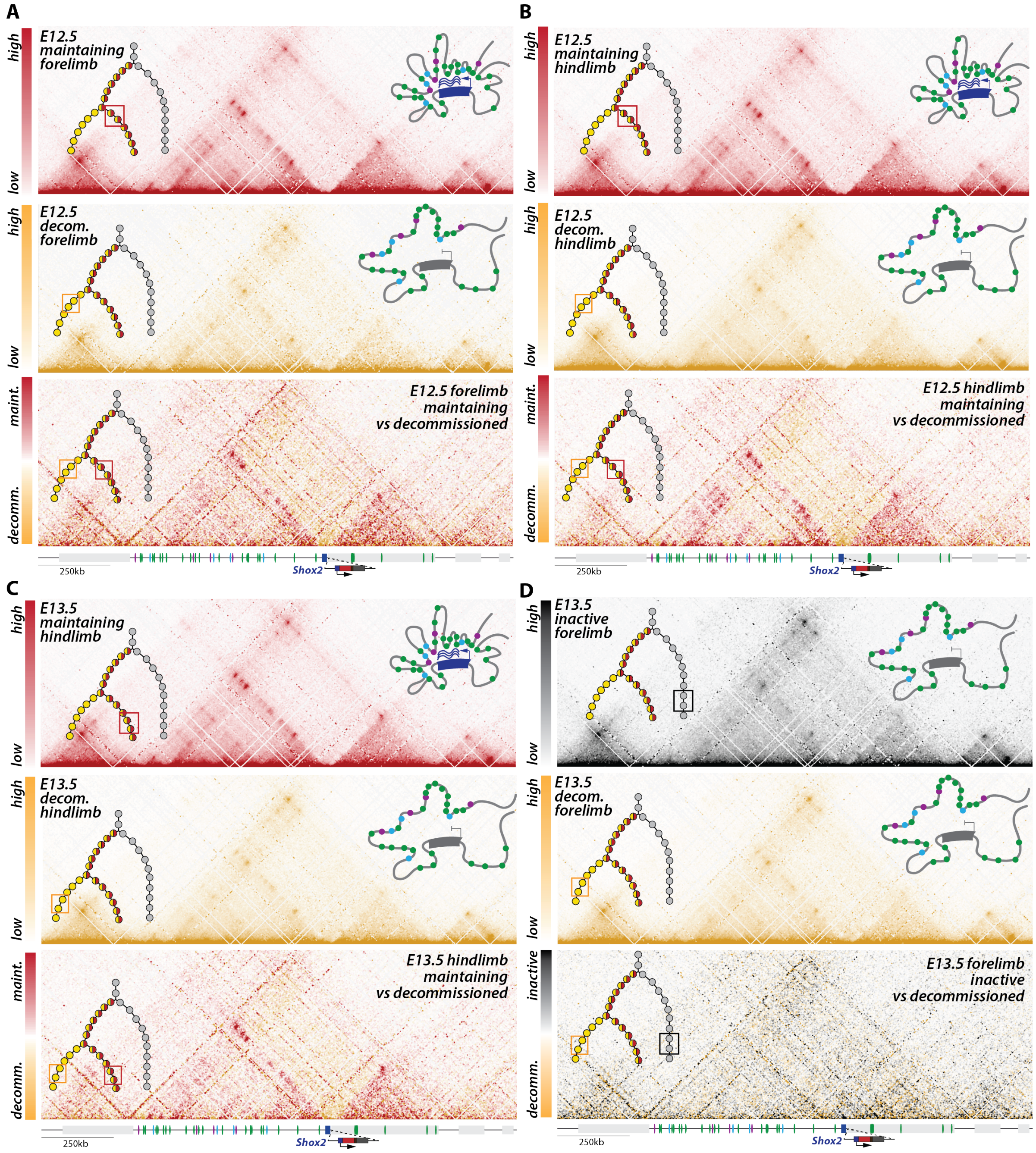


**Supplementary Figure S8.** *In each panel: the upper left illustration represents the position of the investigated cells in the regulatory trajectory; the upper right one a model of the average 3D locus structure. Maps coordinates mm39: chr3:65,781,633-67,435,852. Maint. = maintaining; decomm. = decommissioned. A-C: C-HiC maps of FACS-sorted maintaining cells (top), decommissioned cells (middle) and subtraction map between both (bottom) in* ***A.*** *E12.5 forelimb,* ***B.*** *E12.5 hindlimb and* ***C.*** *E13.5 hindlimb.* ***D.*** *C-HiC maps of FACS-sorted hindlimb E12.5 maintaining cells (top), decommissioned cells (middle), and subtraction map between both (bottom).*

### Supplementary Figure S9

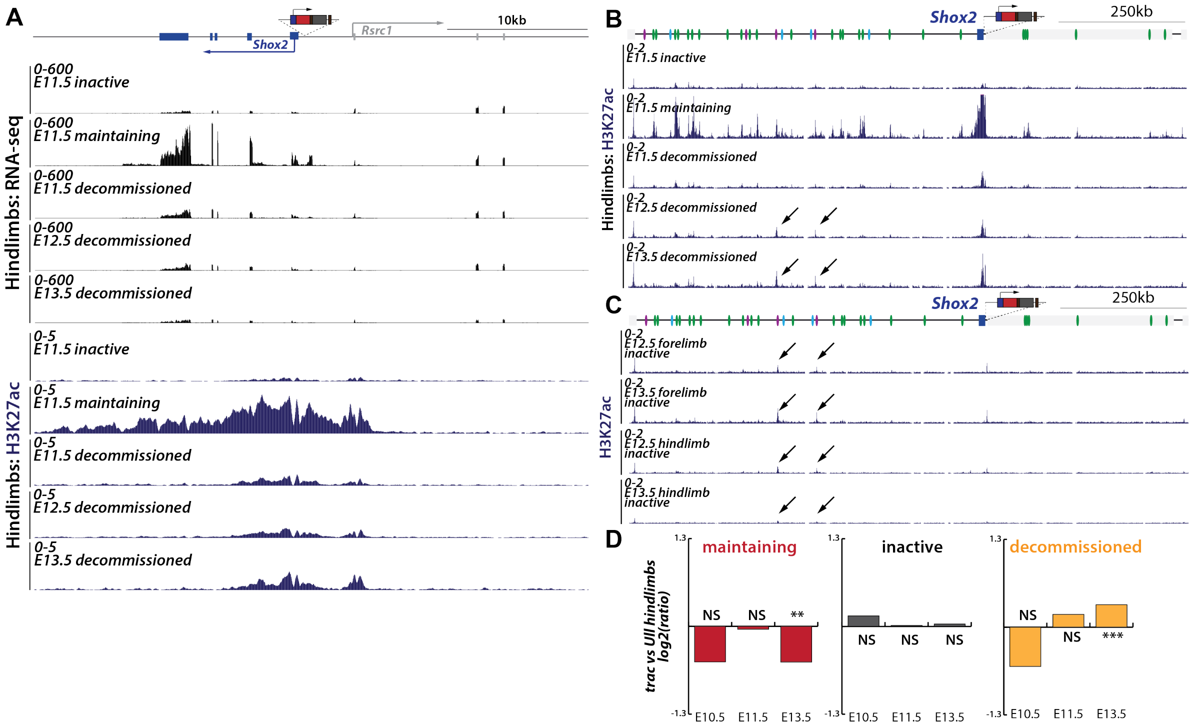


**Supplementary Figure S9:** ***A.*** *RNA-seq, H3K27ac ChIP-seq tracks in early hindlimb inactive, maintaining, decommissioned and late decommissioned FACS-sorted cells at the Shox2 and Rsrc1 promoter regions (mm39: chr3:66,870,000-66,910,000).* ***B.*** *Hindlimb H3K27ac ChIP-seq tracks in early inactive, maintaining, decommissioned and late decommissioned FACS-sorted cells over the Shox2 regulatory landscape (mm39: chr3:66,190,000-67,290,000). Note the loss of H3K27ac at enhancers in decommissioned cells. Note that two of the fours late enhancers show activity in decommissioned cells (black arrows).* ***C.*** *Fore and hindlimb H3K27ac ChIP-seq tracks in early and late inactive cells over the Shox2 regulatory landscape (mm39: chr3:66,190,000-67,290,000). Note that two of the four late enhancers show activity in inactive cells (black arrows).* ***D.*** *Log2 ratio between the proportion of hindlimb Shox2$^{trac}$ and Shox2$^{Ull}$ initiating, maintaining and decommissioned cell populations, identified by flow cytometry analysis, at E10.5, E11.5 and E13.5. T-tests were utilized to calculate p-values from replicates (**See supplementary Figure S10**). NS= non-significant, *=p<0.05, **=p<0.01 and ***=p<0.001.*

### Supplementary Figure S10

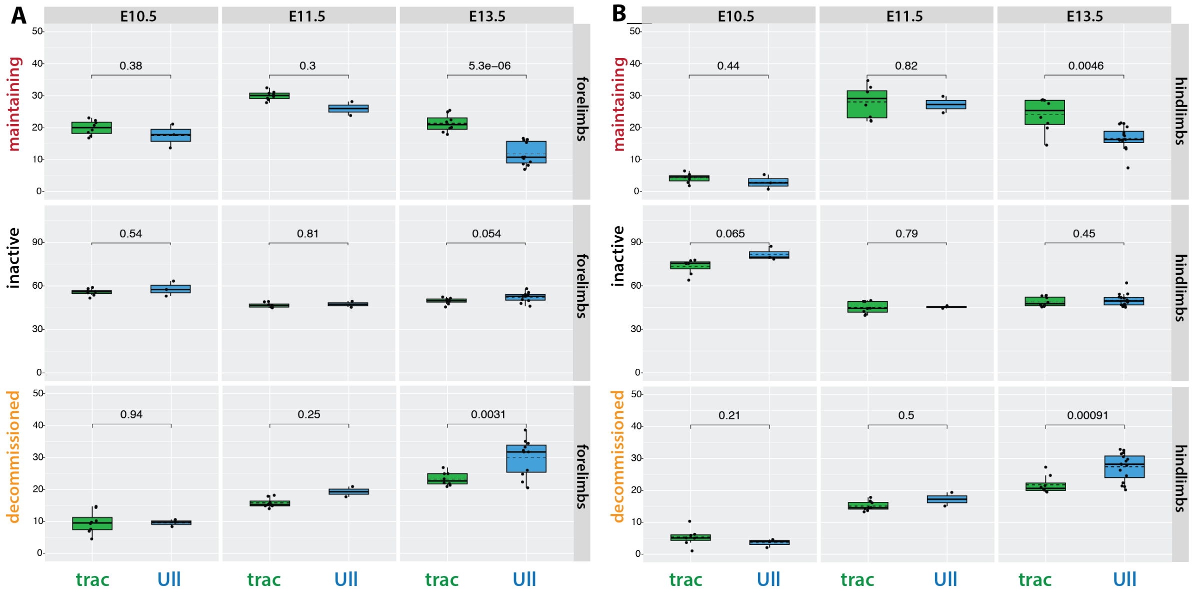


**Supplementary Figure S10.** ***A.*** *Boxplot representation of maintaining, inactive, and decommissioned cell populations, identified by flow cytometry analysis, in Shox2$^{trac}$ versus Shox2$^{Ull}$ forelimbs at E10.5, E11.5 and E13.5. Statistical significance was assessed using T-tests on replicates. Each dot represents one replicate.* ***B.*** *Boxplot representation of maintaining, inactive, and decommissioned cell populations, identified by flow cytometry analysis, in Shox2$^{trac}$ versus Shox2$^{Ull}$ hindlimbs at E11.5, and E13.5. Statistical significance was assessed using T-tests on replicates. Each dot represents one replicate.* ***A-B****, number of replicates for each forelimb genotype: Shox2$^{trac}$ N at E10.5= 8, N at E11.5 = 8, N at E13.5 = 8; Shox2$^{Ull}$ N at E10.5= 3, N at E11.5 = 2, N at E13.5 = 11; and for each hindlimb genotype: Shox2$^{trac}$ N at E10.5= 7, N at E11.5 =8, N at E13.5 = 8; Shox2$^{Ull}$ N at E10.5= 3, N at E11.5 = 2, N at E13.5 = 16.*

**Supplementary Figure S11**

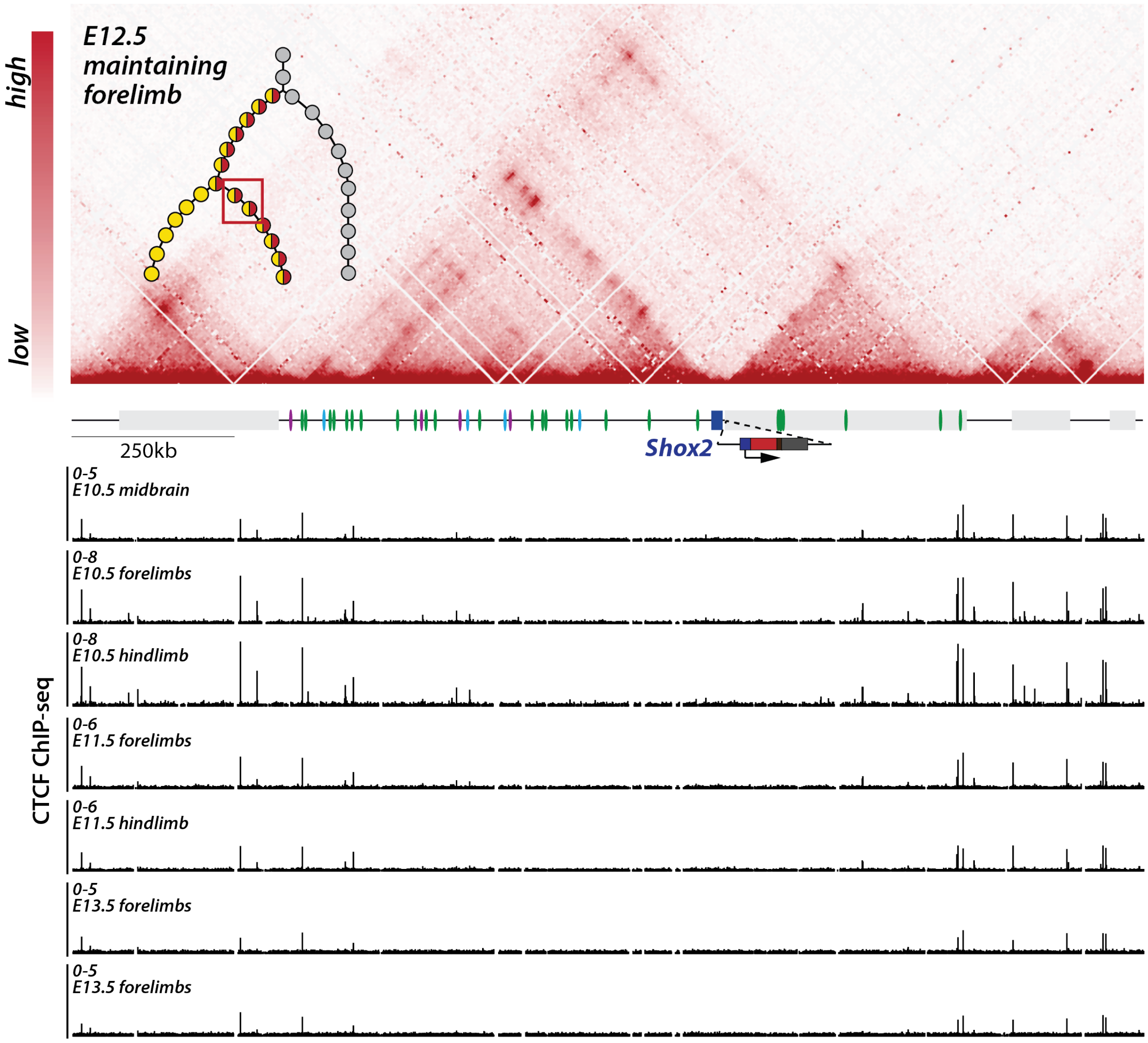


**Supplementary Figure S11. Binding of CTCF at the *Shox2* locus.** *Above: C-HiC map of E12.5 FACS-sorted maintaining forelimb cells. Below: CTCF ChIP-seq tracks of E10.5 midbrain, E10.5, E11.5 and E13.5 fore and hindlimbs (Andrey et al., 2017). Maps coordinates mm39: chr3:65,781,633-67,435,852.*
